# Validation of the host fish relationship between Hickorynut (*Obovaria olivaria*) and Lake Sturgeon (*Acipenser fulvescens*) from wild-caught and archival museum fishes

**DOI:** 10.64898/2026.11.13.764379

**Authors:** Katriina L. Ilves, André L. Martel, Tim Haxton, Sarah E. Steele, Noel R. Alfonso, Benjamin R. Aubrey, Anita LeBaron, Elise Millar, Roger D. Bull, Shauna Fulton, Stéphanie Tessier, Annie Paquet, Marie-Hélène Fraser, Hans-Frédéric Ellefsen

**Affiliations:** Canadian Museum of Nature, Research & Collections, Zoology, Ottawa, ON, Canada; Ministry of Natural Resources, Peterborough, ON, Canada; Ministry of the Environment, Conservation and Parks, Peterborough, ON, Canada; Canadian Museum of Nature, Research & Collections, National Biodiversity Cryobank of Canada, Ottawa, ON, Canada; Ministère de l’Environnement, de la Lutte contre les changements climatiques, de la Protection de la Biodiversité, de la Faune et des Parcs, Ǫuébec, ǪC, Canada; Ministère de l’Environnement, de la Lutte contre les changements climatiques, de la Protection de la Biodiversité, de la Faune et des Parcs, Longueuil, ǪC, Canada; Fisheries and Oceans Canada, Institut Maurice Lamontagne, Mont-Joli, ǪC, Canada

**Keywords:** Freshwater mussels, Unionidae, glochidia, host fish, Acipenseridae, natural infestation, natural transformation, natural history collections, DNA barcoding, larval morphology

## Abstract

Understanding the obligate relationship between freshwater unionid mussels and the host fish(es) of their larvae (glochidia) is key to the conservation of this generally threatened fauna. A host relationship between at-risk species Hickorynut (*Obovaria olivaria*) and Lake Sturgeon (*Acipenser fulvescens*) in Canada has long been presumed based on evidence from laboratory experiments in the United States. This study aimed to make a definitive link in the wild between these two species, while also examining other potential fish-mussel connections in the Ottawa and St. Lawrence Rivers. Using a combination of field collections, morphological and genetic examinations of glochidia from wild-caught fishes, and morphological examination of glochidia from archival specimens from natural history collections, we make a definitive link between these two species for the first time in nature in Canada, with evidence for both natural infestation and natural transformation. This work provides novel methodologies that are widely applicable to studies of mussel-fish relationships and reveals the utility of natural history museum, and other, fish collections for their study.

**Graphical abstract:** 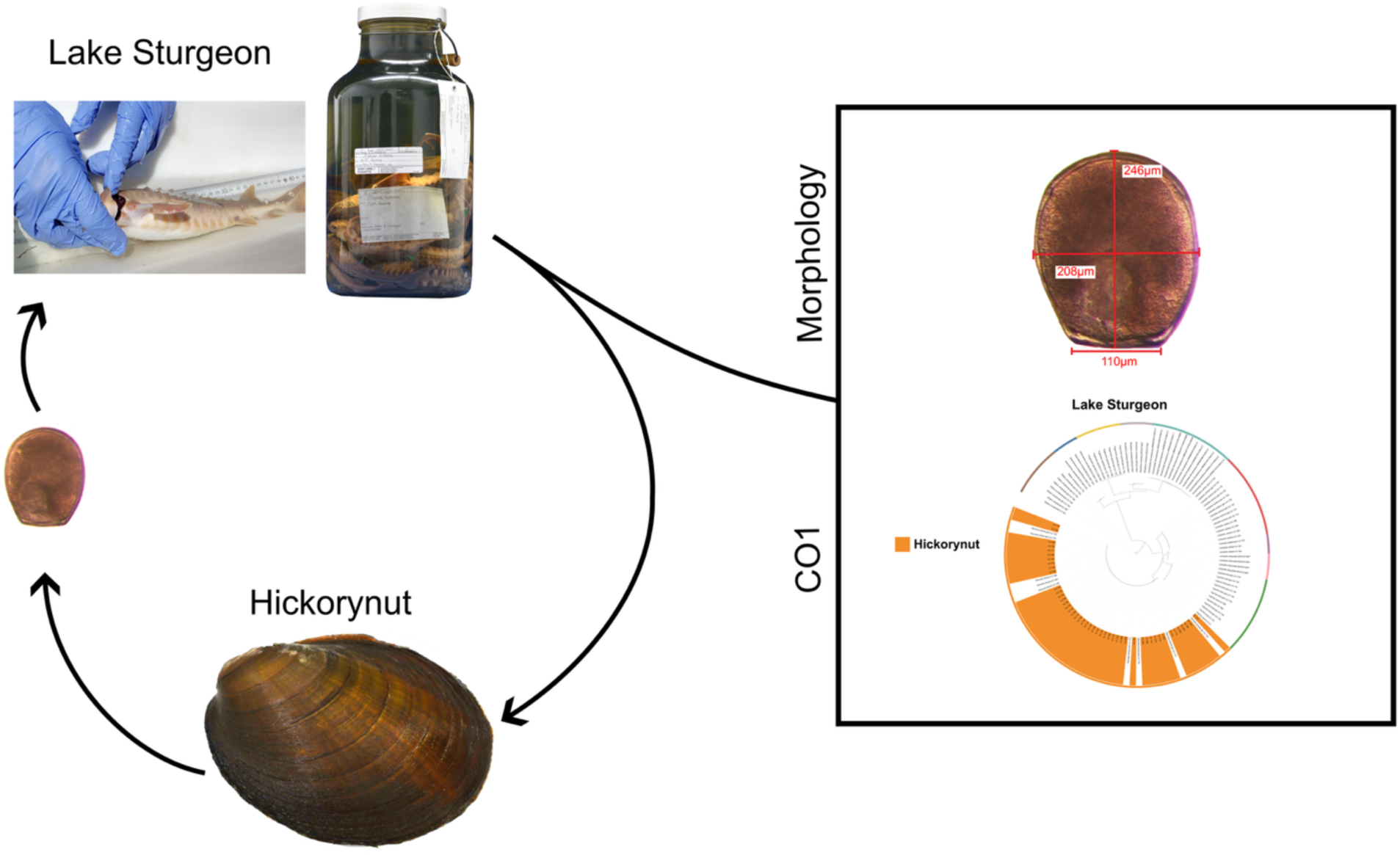

## Introduction

Freshwater mussels (order Unionida) are globally among the most imperiled faunas, with 51% of extant species at risk of extinction (IUCN 2025). Of Canada’s 55 native species, 19 (∼35%) are considered at-risk (Government of Canada 2026), resulting from numerous, often compounding and confounding, threats including, habitat loss and alteration, degraded water quality, invasive species, and altered access to their larval hosts due to dams or other barriers (e.g., Williams et al. 1993; Watters 1996; Nobles and Yang 2011; Zieritz et al. 2022; Aldridge et al. 2023; Nakamura et al. 2023; Prestes et al. 2024; Cushway et al. 2025). As providers of ecological services such as water filtration, nutrient recycling, substrate oxygenation, shells as habitat, and food for wildlife, freshwater mussels are a critical component of healthy freshwater ecosystems (e.g., Vaughn et al. 2008; Vaughn 2018; Zieritz et al. 2022), being labeled as ‘sentinels’ of water quality and ecosystem health (Ferreira-Rodriguez et al. 2023).

All freshwater mussel species native to Canada (two families: Margaritiferidae and Unionidae) have a fascinating life history wherein they require a vertebrate host, typically a fish, to complete their lifecycle (Cummings and Graf 2015; reviewed in Aubrey et al. 2025), although there are rare occasional exceptions of metamorphosis without parasitism for two species (Lefevre and Curtis 1911; Howard 1914; Allen 1924; Dickinson and Sietman 2008). This obligate relationship presumably evolved because it provided the advantage of upstream dispersal, which comes with the ability to colonize new habitats, for an otherwise relatively sedentary fauna (Kat 1984). Fish-mussel connectivity also comes with ecological costs and evolutionary risks, as the reproductive success, recruitment, and persistence of the mussels are dependent on their host(s) (e.g., Reuling 1919; Howard and Anson 1922; Spooner et al. 2011; Douda et al. 2013; Modesto et al. 2018; Dias et al. 2020). Host specialists, those restricted to a single or small number of hosts, are particularly vulnerable to changes in their host’s distribution or abundance (e.g., Neves 1993, cited in Haag 2012; Spooner et al. 2011; Rock and Österling 2026, but see Modesto et al. 2018). Thus, identifying the host relationships of mussels is a key component to understanding their evolutionary and biogeographic histories and for their conservation (McLain and Ross 2005; Allen et al. 2007; Zanatta and Wilson 2011; Schwalb et al. 2013; Modesto et al. 2018, Keogh et al. 2021; Lopes-Lima et al. 2021; Nogueira et al. 2023).

The parasitic larvae of Canada’s native freshwater mussels are called glochidia (sing., glochidium), which are brooded by female mussels in modified gill pouches called marsupia (Cummings and Graf 2015; Graf and Ó Foighil 2000) and vary in morphological and life history characteristics. Each species has one of two basic types of glochidia, hooked or hookless, where glochidia with hooks typically attach to the fins, scales, or other external structures of their host, while those without hooks generally attach to gills (Howard and Anson 1922; Cummings and Graf 2015). Glochidia remain on their hosts for some length of time (a few weeks to many months, depending on the species, water temperature, and other factors) where they obtain nutrients and undergo a metamorphosis into the juvenile stage, at which point they slough off their host and drop to the substrate, where only a small proportion will survive and grow to the adult stage (Howard and Anson 1922; Kat 1984; Barnhart et al. 2008; Cummings and Graf 2015). Further variation in reproduction across species includes the relative length of time females remain gravid (tachytictic/short-term vs. bradytictic/long-term brooders, frequency of brooding, timing (season) of glochidia release, length of time glochidia remain attached to their host(s), and the number and identity of hosts (e.g., Graf 1997; Graf and Ó Foighil 2000; Barnhart et al. 2008; Cummings and Graf 2015). Furthermore, with their relatively simple morphologies and small size (∼80 – 380 um length; Hoggarth 1999), glochidia can be difficult to distinguish based on physical features, particularly to the species level (e.g., Watters 1994; Zale and Neves 1982; Graf and Ó Foighil 2000).

With all of these variables, identifying the host(s) of each mussel species presents many challenges, and most of the research on this subject has been conducted in controlled laboratory environments (Watters et al. 2009; Haag 2012; Aubrey et al. 2025), where potential host fish(es) are exposed to or manually inoculated with glochidia (e.g., Zale and Neves 1982; Yeager and Saylor 1995; Khym and Layzer 2000; Gray et al. 2002; Johnson et al. 2012; Fritts et al. 2012; Hart et al. 2018; Huber and Geist 2019; Skorupa et al. 2022; Stenske et al. 2026). Following the designations of Hoggarth (1992), many studies have been successful in providing evidence of laboratory infestation (LI), where glochidia successfully attach to a host and the presence of (encysted) glochidia on the gills or fins is confirmed. Others have gone a step further and demonstrated laboratory transformation (LT), where metamorphosis from the larval to juvenile stage occurs, which is typically determined by the presence of live juveniles siphoned from the aquarium bottom (e.g. Freshwater Mussel Host Database 2017; evidence for Canadian mussel hosts reviewed by Aubrey et al. 2025).

Although such laboratory experiments are significant with respect to confirming the physiological compatibility of a mussel species and its possible hosts, there are many confounding factors which may not translate to an ecological relationship in the wild (Watters et al. 2009; Haag 2012; Hove et al. 2012). These factors include, synchronicity and sympatry, two factors that are not always accounted for in laboratory trials (e.g., when mussel and fish stocks are sourced from different geographic locations, or when mussels and fishes do not typically co-occur in the wild; Aubrey et al. 2025). For determining host relationships in nature, the most common evidence type is natural infestation (NI), where glochidia are detected on wild-caught fishes and are identified based on morphological characters, typically three linear measurements (height, length, hinge length; Hoggarth 1988, 1999; Kennedy and Haag 2005) or genetic methods (e.g., DNA barcoding; Kneeland and Rhymer 2007, 2008; Dobbs et al. 2024; Poelmann et al. 2024). Although this points to a potential host relationship by demonstrating there was not an immediate immune rejection by the host, it is not as definitive as evidence of glochidial metamorphosis found on fishes in nature (natural transformation (NT)). Given the complexity of the field and lab work required to demonstrate NT, this is the most challenging and rare study of mussel-host relationship (Watters et al. 2009; Haag 2012; Aubrey et al. 2025). Most studies of NT involve collecting naturally infested fish(es) from the wild and maintaining them in laboratory conditions to monitor the aquaria for living juveniles that have dropped off the fish (e.g., Boyer et al. 2011; Hove et al. 2012; Van Susteren et al. 2015; Sietman et al. 2018; Robinson et al. 2025). Some studies on naturally-infested fishes have instead made histological examinations of glochidia development (e.g., Karna and Millemann 1978; Ieshko et al. 2016), used size change as an indication of metamorphosis (Cunjak and McGladdery 2011), or noted the presence of paired adductor muscles (Hove et al. 2011) to demonstrate larval transformation into a post-metamorphic or juvenile stage.

A host relationship between Hickorynut (*Obovaria olivaria*) and Lake Sturgeon (*Acipenser fulvescens*) in Canada has long been suspected (COSEWIC 2011). Hickorynut is a thick-shelled species that prefers moderate to deep waters and sandy river beds (Parmalee and Bogan 1998) and is distributed in large waterbodies of the Mississippi River watershed and the Great Lakes/St. Lawrence basin (COSEWIC 2011). It is listed as Endangered in Canada (Government of Canada 2019), making confirmation of its host fish(es) of importance for conservation planning. In Canada, its primary host has been presumed to be Lake Sturgeon, based largely on a lab experiment (LT; Brady et al. 2004) and range correlation (i.e., Hickorynut is only found in areas where Lake Sturgeon is present). It has also been speculated that in Canada Atlantic Sturgeon (*Acipenser oxyrinchus*) may also be a suitable host (COSEWIC 2011). To our knowledge, a Hickorynut-Lake Sturgeon relationship has never been demonstrated in nature (Freshwater Mussel Host Database 2017; Aubrey et al. 2025).

Here we aim to validate the presumed host relationship of Hickorynut and Lake Sturgeon in nature by demonstrating both NI and NT using a combined morphology- and genetics-based approach to identifying glochidia and their developmental stage from wild-caught fishes. Globally, sturgeon species are known to be hosts for multiple mussel species (Freshwater Mussel Host Database 2017) and the number of hosts identified for a mussel species is correlated with the number of studies of that species’ host relationships (Hewitt et al. 2019; Aubrey et al. 2025); therefore, we also examined a community of fishes and mussels to assess whether Lake Sturgeon may be a host to multiple mussel species, and/or if other fishes may be hosts for Hickorynut. Finally, we explore the utility of natural history museums and other fish collections as potential sources of evidence for NI for mussel-host relationships. This study’s combined morphology-genetics approach, where *individual* glochidia were identified based on a novel morphological key and sequenced for the mitochondrial DNA barcoding gene cytochrome *c* oxidase subunit I (COI), provides additional characters and methods for identifying Hickorynut glochidia from wild-caught fishes. From this work, we provide the first evidence of both NI and NT of Hickorynut glochidia on wild-caught Lake Sturgeon in Canada. This relationship is further supported by the first published use of museum and government agency fish collections for determining host fish of mussel glochidia.

## Materials and methods

### Taxonomy

For mussels, the common and scientific names follow Harris et al. (2025) and those of fishes follow Page et al. (2023). We acknowledge several published proposals for taxonomic changes in Acipenseridae, and discussion thereof (Brownstein and Near 2025; IUCN Sturgeon Specialist Group 2025; Kottelat and Freyhof 2025; Hilton et al. 2026; Near and Brownstein 2026). These proposals, with others likely to follow, are currently under review by the Joint Committee on Common and Scientific Names of Fishes from the United States, Canada, and Mexico (https://www.asih.org/nfjc), and as such we here use the currently accepted scientific name of Lake Sturgeon as *Acipenser fulvescens*.

### Fieldwork

Ethics approval was obtained from the Canadian Museum of Nature’s Animal Care Committee (Protocol I.D. #: 2021.05.002) before starting the study and permits were obtained from the Ministère des Forêts, de la Faune et des Parcs (MFFP; name of Ministry at time of sampling & permits)/Ministère de l’Environnement, de la Lutte contre les changements climatiques, de la Protection de la Biodiversité, de la Faune et des Parcs (MELCCPBFP; current name of Ministry): 2021 SP3056R03-07-14-16, PM_22-07-SP-002-GR-0, 2023 05 29 021 00 SP, 2023 05 26 020 00 SP, 2023 06 16 023 00 SP, 23-07-SP-002-GR-0, PM_23-07-SP-004-GR-0; and Fisheries and Oceans Canada: MPO LEP ǪC 21011, MPO LEP ǪC 22012, MPO LEP ǪC 23007, MPO LEP ǪC 23024, MPO LEP ǪC 23019.

#### Ottawa River sampling

We sampled the Lac Coulonge region of the Ottawa River in 2021 (September 20-22; November 8-9), 2022 (November 8-11), and 2023 (October 31 – November 2) (Fig. 1A, Table S1). Seining at a few sites was conducted in September 2021 (Finlay Islands) and November 2022 (Woods Island) using a 70 x 4 ft (21.3 m x 1.2 m) seine with 1/8 in (3.175 mm) mesh and a 4 x 4 ft (1.2 x 1.2 m) pocket. The bulk of the sampling was via gillnet. Multi-panel gillnets of varying lengths, heights, and mesh sizes were set for ∼1.5-24 hours (Small: 10.7 m x 0.9 m; 7 mesh sizes x 1.5 m panels: 13, 19, 25, 32, 38, 44, 51 mm; Large: 24.6 x 1.8 m; 8 mesh sizes x 3.1 m panels: 38, 51, 64, 76, 89, 102, 114,127 mm). Nets were set to target juvenile sturgeon based on previous netting studies (Haxton and Friday 2020) and depth preferences (Haxton 2011).

**Figure 1.**
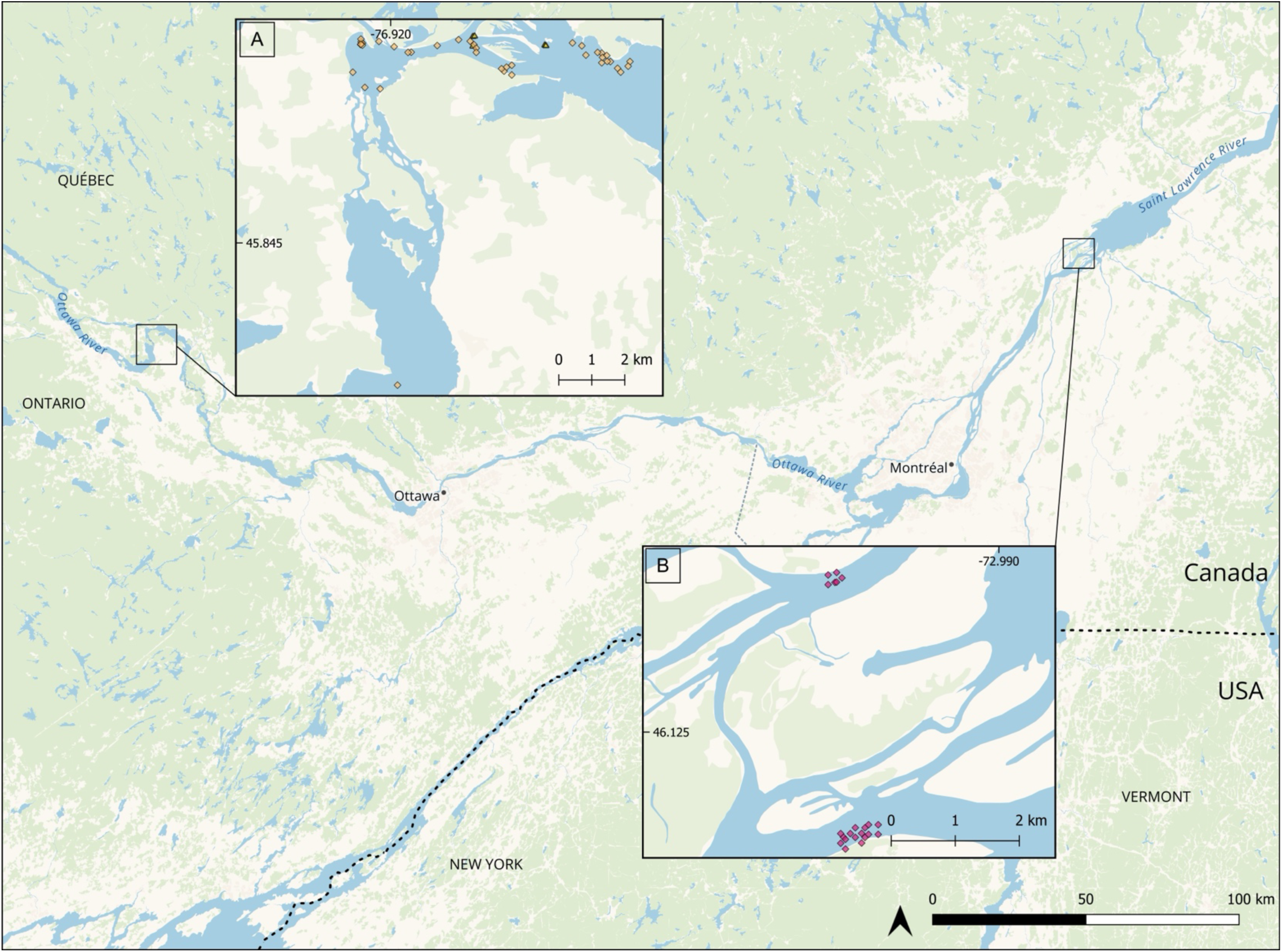
Map showing sample sites in the (A) Lac Coulonge region of the Ottawa River and (B) Lac Saint-Pierre archipelago region of the St. Lawrence River. Sample codes and coordinates can be found in Table S1.

#### St. Lawrence River sampling

The Lac Saint-Pierre archipelago was selected as a study site because of its high abundance of juvenile Lake Sturgeon and the opportunity to examine individuals during the ongoing biennal recruitment monitoring program. In addition, numerous Hickorynuts were incidentally captured in gillnets in 2019 (Fraser and Lamoureux 2021). A diving survey conducted in 2020 further confirmed the abundance of Hickorynuts in this section of the St. Lawrence River (MELCCPBFP and DFO, unpublished data).

On June 15 and 16, July 20, and August 17, 2021, we joined a team from the MFFP/MELCCPBFP during their biennial Lake Sturgeon recruitment monitoring program in the Lac Sainte-Pierre archipelago of the St. Lawrence River (Fig. 1B, Table S1). Multi-panel gillnets (45.6 x 1.8 m) with mesh sizes 25, 38, 51, 64, 76, and 102 mm were deployed perpendicular to the shoreline at depth ranging from 4 to 10 m. Nets were typically set for approximately 24 hours (∼21-27 hours), although a few shorter sets (1.5-5.75 hours) were also conducted. After the MFFP/MELCCPBFP team had completed their sampling procedure aboard the primary vessel, individual Lake Sturgeon were transferred to a second boat, where they were measured (total length), and their gills were examined for the presence of glochidia following the protocol described below. Fin clips were collected from a subset of individuals and preserved in 95% ethanol (EtOH) for subsequent genetic analyses. Due to the large number of Lake Sturgeon captured, this was the only fish species examined for glochidia in the field. Some deceased bycatch was retained and taken back to the CMN laboratory for later examination. Additional sampling was attempted in the same area from May 16-18, 2023; however, unusually high-water levels and strong currents associated with a late spring limited sampling efficiency. As a result, only eight Lake Sturgeon were captured over three days and none showed evidence of glochidia.

#### Examination of live fishes for evidence of glochidia on gills

To minimize the number of fishes sacrificed, we examined nearly all individuals live in the field and only humanely euthanized those specimens where there was evidence of glochidia on their gills. Such evidence included the presence of obvious glochidia (larval shell or hinge visible), cysts, or other aberrations on a gill filament of the appropriate size that could not be dislodged with a stream of water from a squirt bottle and that could potentially mask a glochidia lying underneath some gill tissue. Images of the field setup and materials can be found in the Supplementary material (Fig. S1). The gill examination procedure generally involved at least three people and for large or active fish, a fourth person was sometimes needed to safely restrain the fish. We used a V-shaped PVC measuring trough that was propped up on one end with rocks or other material to hold water at the lower end to keep one set of gills hydrated as the other set was examined. The fish for examination was placed on its right side, its total length measured (mm), and its left gills examined for evidence of glochidia by first gently lifting the operculum. One person held an LED light to illuminate the gill arch under examination and another member of the team (KLI, NRA, AP, HFE, or MHF at the St. Lawrence River sites and usually ALM at the Ottawa River sites) used a 4, 8 or 10x hand lens to check the gill filaments for possible evidence of glochidia. One to two additional people were required to hold the fish steady in the trough. Water was regularly streamed across the gills using a squirt bottle to both keep the gills hydrated and to dislodge occasional gill mucus and debris. This examination procedure was repeated for the inner and outer face of each gill arch by gently lifting the arch with a smooth-edged metal spatula, taking care not to damage the gill filaments. Once the left side of the fish was examined, the fish was turned over, and the examination was repeated on the right side. For large active fishes (e.g., Northern Pike (*Esox lucius*) >750 mm), examination took place with the fish in a large bin rather than in the measuring trough (e.g., Fig. S1). A fin clip was taken from most specimens and preserved in 95% EtOH. If examination of a single fish took an extended time (> 5 min) or the fish showed distress by vigorous body or opercular movement, it was placed in a recovery bin with fresh river water until the behaviour subsided and the examination could recommence. After examination, fishes with no evidence of glochidia were kept in a large recovery bin with fresh water for at least 20 minutes before being released. Fishes with possible evidence of glochidia were humanely euthanized using MS-222 following the approved procedures in our Animal Use Protocol. If time and weather conditions permitted, gill arches from euthanized fishes were dissected in the field and preserved in 95% EtOH (in a jar or Falcon tube) as this was the best way to rapidly preserve the glochidial DNA. Otherwise, the entire fish was frozen, and the gills were later extracted and gill arches separated into individual glass vials in 95% EtOH in a laboratory at the CMN’s Natural Heritage Campus.

### Labwork

#### Control samples for morphological and genetic identification of glochidia

A control data set consisting of the seven freshwater mussel species found in the Lac Coulonge region of the Ottawa River was compiled for both morphological and genetic characterization of glochidia. For those controls, gravid female mussels, from which mature glochidia and adult tissues could be extracted and analysed in the laboratory afterwards, were collected during snorkeling and SCUBA diving in the Ottawa River. Gravid female mussels collected included Triangle Floater (*Alasmidonta undulata*), Eastern Elliptio (*Elliptio complanata*), Plain Pocketbook (*Lampsilis cardium*), Eastern Lampmussel (*Lampsilis radiata*), Black Sandshell (*Ligumia recta*), Hickorynut, Giant Floater (*Pyganodon grandis*), with an additional three species, Cylindrical Papershell (*Anodontoides ferussacianus*), Fatmucket (*Lampsilis siliquoidea*), and Creeper (*Strophitus undulatus*), included for genetic analyses only. These latter three species have not been documented by ALM in the immediate region but have been found in upstream and tributary populations that could possibly send glochidia to the Lac Coulonge area and iNaturalist records suggest they are in or nearby the study region. Details of the voucher (or other) specimens and the associated accession numbers for the molecular control dataset are in Table S2, while the details and measurements of the adult vouchers and glochidia are in Tables S3 and S4, respectively.

#### Molecular identification of glochidia

We targeted the mitochondrial COI gene, commonly used for species-level DNA barcoding in molluscs and other animals (Hebert et al. 2003; Campbell et al. 2008; Layton et al. 2014; Robinson et al. 2025). Control sequences of 10 species were mostly generated in this study, except for those of Cylindrical Papershell, Fatmucket, and Creeper, which were obtained from the Barcode of Life Database (BOLD; Ratnasingham et al. 2024) or GenBank (Benson et al. 2012). Sequences from adults and glochidia were all obtained from specimens sequenced in the CMN’s Laboratory of Molecular Biodiversity (Table S2) following the extraction, PCR, and sequencing protocols detailed below.

#### Processing glochidia from female marsupia and fish gills

All morphological measurements of glochidia were made using microtools and other specialized vessels suitable for manipulating the small organisms, many custom-made (Fig. S2). These tools included entomology forceps, thin microneedles (made of a 100, or 150, or 300 µm diameter insect pin mounted on a fine 2-mm diameter wooden handle), microscissors (for cutting fish gill filaments), glass petri dishes with tiny circular depressions or troughs to immobilize glochidia. Natrolite crystals placed inside the tiny depressions or troughs of the glass dish were commonly used for propping up and immobilizing glochidia in a vertical orientation needed for depth measurements and taking hinge characterization photos.

For examination, a gill arch was placed in a small glass petri dish, and the inner and outer face of each arch was thoroughly examined for the presence of glochidia using Olympus SZX12 and Olympus SZX16 stereo microscopes. If the presence of a glochidium was positively confirmed and was only going to be characterized genetically, the entire excised fragment – fish gill filament and glochidium – was placed in DNA extraction buffer and the DNA extraction process described below was started immediately. If it was first to be used for morphological study, the filament was cut below the glochidium using microscissors and as much fish tissue as possible was removed through careful manipulation of microneedles to tease the tissue off without cracking the prodissoconch (larval shell). Once sufficient tissue was removed to take accurate measurements, the glochidium was transferred to a smaller glass petri dish with small depressions and a shallow solution of 30-50 % EtOH and natrolite crystals. Measurements were taken as described below and if the glochidium was to also be characterized genetically, it was transferred via micropipette into a 1.5 mL Eppendorf tube containing DNA extraction buffer and the DNA extraction process was started immediately.

#### Examination of fishes from museum and provincial archival collections

Archival specimens of Lake Sturgeon from the CMN’s Fish Collection and from the MELCCPBFP’s facility in Longueuil, ǪC (Table S5) were examined in the lab for presence of glochidia by first examining each gill face under a dissecting microscope. If a possible glochidium was located, the gill filament was excised for further examination following the procedure outlined above (section *Processing glochidia from female marsupia and fish gills*).

#### DNA extraction: glochidia

Due to the small size of the glochidia, we used an alkaline DNA extraction method, modified from Rudbeck and Dissing (1998) and Malagó et al. (2002), that was designed for forensic work on small amounts of biological material. Glochidia were either pipetted (those used for morphological measurements or taken from female marsupia as controls) or placed with forceps (those directly from gill filaments, not used for morphological measurements) into a 1.5 mL Eppendorf tube containing 20 µL of 0.2 M NaOH, immediately incubated at 75 °C for 20 minutes with shaking at 500 rpm in an Eppendorf ThermoMixer C, after which the reaction was neutralized by the addition of 180 µL of 0.04 M Tris-HCl, pH 7.5. DNA extracts were stored at 4 °C (short-term, up to 7 days), transferred to -20 °C (mid-term, 1-8 weeks), and ultimately stored at -80 °C.

#### DNA extraction: adult controls

DNA was extracted from a small piece of mantle tissue (∼2 mm^3^) using a protocol modified from Ivanova et al. (2006). Recipes for buffers described below are in the Supplementary Material. An incubation buffer consisting of 180 µL Animal Lysis (AL) buffer and 6 µL Proteinase K (20 mg/mL) per sample was prepared and added to the tissue samples in 1.5 mL Eppendorf tubes. The tubes were vortexed for 15 s and placed in a rotating rack in a 56 °C incubator for 1-24 hours to complete lysis (no visible tissue). The tubes were again vortexed for 15 s, 200 µL AMB Buffer was added followed by 200 µL 99% EtOH, with vortexing following each reagent addition. The contents of each tube were then pipetted into a separate silica membrane spin column-collection tube combination (EconoSpin, Epoch Biolabs 1920-250) and centrifuged at 8,000 rpm for 1 min in an Eppendorf 5424 Centrifuge. The flow-through was discarded and 500 µL of First Wash Buffer (150 µL AMB Buffer + 350 µL 99% EtOH) was added to each sample spin column, followed by centrifugation at 8,000 rpm for 1 min. The flow-through was discarded and 500 µL of Second Wash Buffer was added to each sample spin column and centrifuged at 14,000 rpm for 3 min. The flow-through was discarded and the spin column was transferred to a sterile 1.5 mL Eppendorf tube for the final elution steps. A double-elution step was completed with each step including the addition of 75 µL of warmed (56 °C) THE Buffer added directly to the column membrane, incubation at room temperature for 5 min, followed by centrifugation at 15,000 rpm for 1 min. The spin column was discarded and the ∼150 µL DNA extract was stored at 4 °C (short-term, up to 7 days), transferred to -20 °C (mid-term, 1-8 weeks), and ultimately stored at -80 °C.

#### Primer design and nested PCR

PCR on the alkaline extractions was usually completed within one week of extraction. Because we were targeting single very small glochidia, amplification of COI was completed in a two-stage nested PCR protocol. The first stage used primers from Campbell et al. (2008), UnionidCOIF 5’-GTTCCACAAATCATAAGGATATTGG-3’ and UnionidCOIR 5’ TACACCTCAGGGTGACCAAAAAACCA-3’ with the following reaction components in a 15 µL reaction: 1 µL DNA extract, 11.3 µL dH_2_0, 1.5 µL ThermoFisher 10X DreamTaq Buffer, 0.3 µL 10 mM dNTP mix, 0.375 µL of each primer (10 µM), and 0.15 µL ThermoFisher DreamTaq polymerase. The second stage of the nested PCR used one of two internal primer pairs designed for this study using Primer3 implemented in Geneious from an alignment of *Lampsilis* and *Obovaria* COI sequences: COIF-INT1 5’-TGTTRGCGTTRTGGTCTGGT-3’ and COIR-INT1 5’-AGCACCAGCCAAAACAGGYA-3’ or CO1F-INT2 5’-TTRATTCGGGCTGARTTGGG-3’ and COIR-INT2 5’-AACAGCCCAYACAAACAACG-3’. The reaction conditions were the same as for the first stage PCR, except that PCR product from the first PCR was used instead of DNA as the template. The PCR product was often diluted 1:5 or 1:10 before use in the step 2 PCR (e.g., if smearing was noticed on a gel of PCR2 product, PCR2 was rerun with a dilution series of PCR1). The Eppendorf Mastercycler pro S thermal cycler conditions were the same for both PCR 1 and 2, with an initial denaturing step at 95 °C for 3 min, followed by 35 cycles of 95 °C for 30 s, 48 °C for 30 s, and 72 °C for 1 min 30 s and a final extension step of 72 °C for 10 min. A negative control was included for each primer combination at each PCR step. PCR products after PCR2 were visualized on a 1.5 % agarose gel (BioShop agarose + 1x TAE Buffer stained with 0.25 mg/mL ethidium bromide). Two µL of PCR product was combined with 4 µL of 3X Bromophenol Blue Loading Buffer, along with a well containing 4 µL GeneRuler 1 kb DNA Ladder Plus (Fermentas) and visualized with a Ǫuantum ST5 Gel Documentation System (Montreal Biotech Inc.). If a band appeared in the negative control (rare), the PCR reagents were replaced and the process redone.

#### Sanger sequencing

Successful PCRs (bands in the expected range of around 600 bp) were diluted with either 0, 15, 30, or 60 µL of sterile-filtered H_2_O depending on their relative brightness on the agarose gel (i.e., relative concentration) after which a 10 µL sequencing reaction in a 96-well plate for each sample and primer (forward & reverse) was completed with the following conditions: 1 µL PCR product, 0.5 µL BigDye, 0.5 µL 10 µM primer, 1.8 µL 5x ABI Buffer, and 6.2 µL dH_2_O. The sequencing reaction included an initial 3 min denaturation at 95 °C followed by 30 cycles of 96 °C for 30 s, 50 °C for 20 s, and 60 °C for 4 min. For sequencing reaction purification, 1 µL EDTA (125 mM), 1 µL NaOAc (3 M), and 25 µL EtOH were added to each well of an ABI plate and it was spun in an Eppendorf 5804 R Centrifuge at 3700 rpm for 30 s. The full 10 µL of the sequencing PCR reaction was then added to the respective well of the ABI plate and incubated at room temperature away from light for 20 min, after which it was spun at 4 °C for 45 min at 1650 x g. The plate was then inverted onto paper towel within the centrifuge plate holder and spun for 30 s at 180 x g. The plate was then set upright, 35 µL of 70% EtOH was added to each well, and spun at 4 °C for 15 min at 1650 x g. The plate was again inverted onto paper towel and spun for 1 min at 180 x g after which it was dried in a vacufuge (covered from light) for 15 min. In a fume hood, 15 µL of HIDI formamide was added to each well to resuspend the dry DNA pellet and it was spun for 1 min at 3700 rpm. Sequences of purified reactions were visualised via automated capillary electrophoresis on an Applied Biosystems 3500xL Genetic Analyzer (ThermoFisher Scientific).

#### Sequence alignment and phylogenetic analyses

Control sequences of the 10 mussel species potentially distributed in the Lac Coulonge region were obtained from extraction, PCR, and sequencing of adult and/or glochidia from adult female marsupia of expert-identified (ALM or AP) mussels, from BOLD, or GenBank (Table S2).

Sequence editing and alignment were conducted in Geneious Prime 2025.2.1, or earlier (https://www.geneious.com) by KLI. After import, sequences were manually trimmed at the 5’ and 3’ ends and forward and reverse sequences (as available) were aligned using the pairwise alignment option in MUSCLE 5.1 (Edgar 2022) with default settings. Following alignment, any base call with a quality level below 20 was converted to an “N”. Consensus sequences for pairwise (or multiple if sequences were generated from both pairs of internal primers) alignments were generated with default settings. Control sequences were aligned separately.

COI consensus sequences from glochidia from each of the host fish species were aligned with the COI control sequences alignment using the ClustalOmega (Sievers and Higgins 2017) option in Geneious Prime. Phylogenetic trees of the glochidia from each host fish were estimated with IǪ-TREE 3.0-2.1 (Wong et al. 2026), with reduction of severe model violation SH-aLRT test (Guindon et al. 2010), model selection using ModelFinder (Kalyaanamoorthy et al. 2017), and branch supports from the ultra-fast bootstrap method (Hoang et al., 2018). Species identity of the unknown glochidia was determined based on monophyletic grouping of the sequences within the clades of the control sequences.

Phylogenetic trees were mid-point rooted and visualized using Interactive Tree of Life (iTOL) v7.5 (Letunik and Bork 2024).

#### Morphological characterization of glochidia from gravid females with verified species identity

All observations, images, and measurements of glochidia were taken with an Olympus SZX16 stereo microscope, Olympus SDF PLAPO 1XPF and 2XPF lenses, Olympus DP28 Camera with Olympus UTV0.63X C-mount adapter, and cellSens Entry software (https://evidentscientific.com/en/products/software/cellsens) by ALM. When possible, four linear measurements were taken of each glochidium: three standard measurements of height, length, hinge length, and a fourth measurement of depth/width (Fig. 2A). The presence or absence of hooks on the ventral margin of the glochidium valves was noted (Fig. 2B). A qualitative hinge character was also noted about the relative strength of a “whisker-like” structure: Type 1: no structure, Type 2: weak structure, Type 3: strong structure (Fig. 2C).

**Figure 2:**
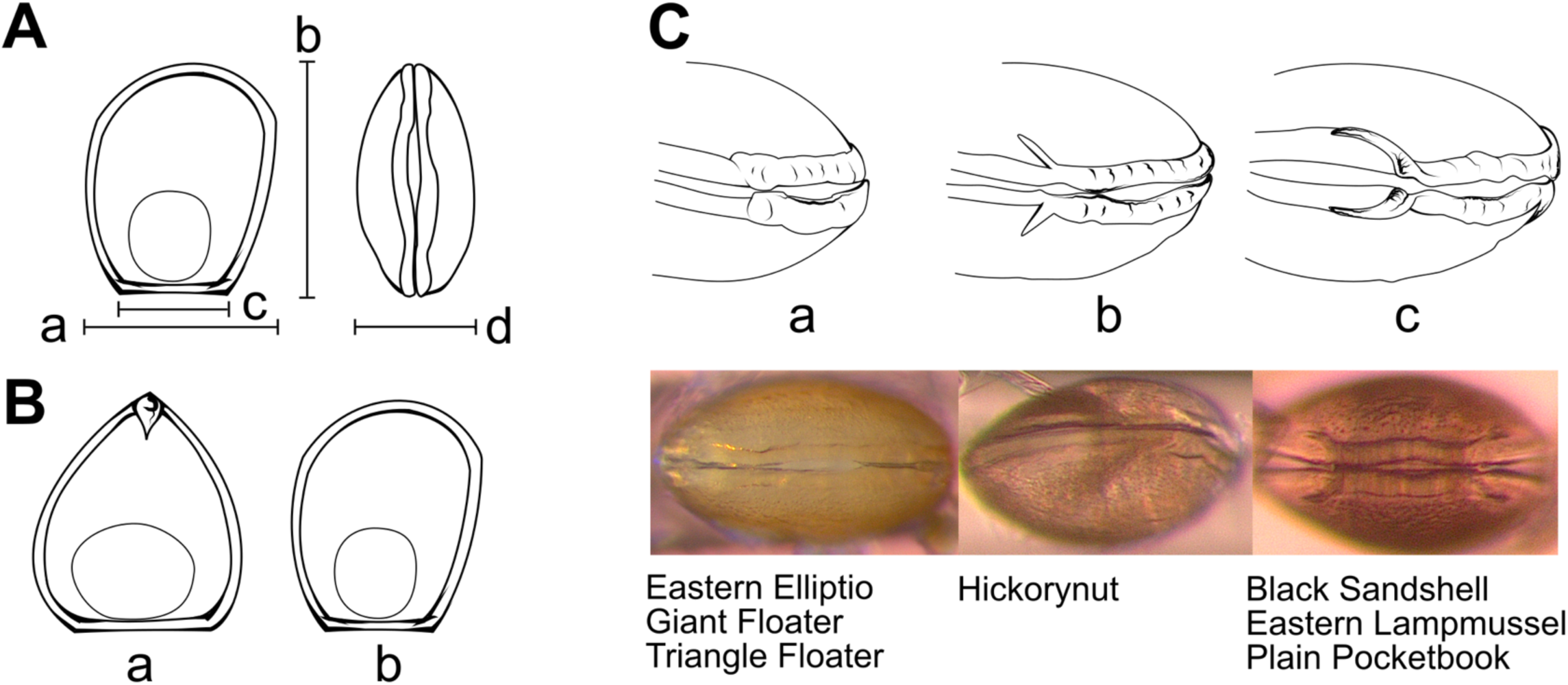
Linear measurements and morphological characteristics used for the morphological species identification of glochidia. Panel A: Linear measurements: Length (a), Height (b), Hinge Length (c), and Depth (d). Line drawing representative of Plain Pocketbook (*Lampsilis cardium*); Panel B: Presence (a) or absence (b) of hooks on the ventral valves. Line drawing representative of (a) Giant Floater (*Pyganodon grandis*) and (b) Plain Pocketbook; Panel C: Whisker character on the hinge: No structure (a, Type 1), Weak structure (b, Type 2), Strong structure (c, Type 3). Line drawing and photos representative of (a) Giant Floater, (b) Hickorynut (*Obovaria olivaria*), and (c) Plain Pocketbook. Species exhibiting each Whisker Type are listed below their respective photos. Note: The orientation of the line drawing C (b) differs from that of C (a,c) as Hickorynut only displays a whisker structure on the anterior end. Figure edited with Inkscape (Inkscape Project (2020) Inkscape (Version 0.92.5). Retrieved from https://inkscape.org)

Glochidia were extracted from the marsupial pouches of gravid female mussels of known species identity (expert identification by ALM or AP) using microforceps and placed in a depression-containing glass petri dish with a solution of 30-50 % EtOH supplemented with natrolite crystals. Under the microscope with magnification 7-115X (using 10X eyepieces), microforceps and microneedles were used to position the glochidium in the correct orientation to take the four linear measurements of the prodissoconch – in plane for height, length, hinge length, and vertical/perpendicular for depth/width.

All statistical analyses were conducted using R 4.5.3 (R Core Team, 2026). 95% confidence intervals, minimum, and maximum were calculated for each linear measurement for each species to define a range of expected values for identification of gill-extracted glochidia (test data). For the lower and upper bounds, the smallest and largest values calculated were used for the expected range. This provided a conservative estimate of all possible values likely to be observed in the test data. Visualization of variation within and among species of the stock (training) data was conducted with Principal Component Analysis (prcomp) and 3D plotting (scatterplot3d, Ligges and Mächler 2003; rgl, Murdoch and Adler 2026). Species polygons were created using 95% confidence intervals to assign shape space to one or multiple species.

#### Gill (Test) Dataset: Glochidia extracted from gills of host fishes – species identity unknown

Glochidia were extracted from host gills, imaged, and measured (ALM) for the same four linear measures, presence or absence of hooks noted, and the hinge corner imaged for whisker characterization (SES/ALM). Linear measurements were then individually compared to the expected ranges calculated from the stock data to classify each measurement by one or multiple species identities. Likewise, the whisker images (if available) were examined and classified by one or multiple species identities. Multiple species identities were labelled as “Ambiguous” for visualization purposes and further analysis. If the observed value did not fall within the expected range of any species found in the stock data, the observation was classified as “Unknown”. For each glochidium, the four linear species identity classifications, one hook identity classification, and one whisker identity classification were compared for congruence. If any classification was a single species, the glochidium was only identified as that species if all other measurements, where available, supported the identification (either as a single species identification, or that species appeared within the group of ambiguous identifications). If there was no congruence among the classifications, the species identification remained “Ambiguous”. If any measurements contained an “Unknown” classification, the species identification remained “Unknown” unless another classification of a single species with absolute certainty (no overlap with any other species) was made. As our statistical and visualization methods could not handle N/As, glochidia without all four linear measurements were excluded from analyses. Species assignments were then visually compared with the stock data visualizations and species polygons to check for congruence. PCA was conducted on the pooled training and test data so that the datapoints were aligned in the same spaces allowing for direct comparisons visually. Finally, Linear Discriminant Analysis (LDA) was performed to see how well the manual species assignments performed compared to a multivariate statistical approach. The training and test datasets were centered and scaled. Since hinge length was found to be highly correlated with length (r = 0.96), it was removed from the analysis. The LDA was performed using the ‘lda()’ function in the MASS package (Venables and Ripley 2002), and predictions of species identification of the glochidia from the gills (training data) were calculated. A confusion matrix was produced to investigate discrepancies between the manual identification of the gill glochidia and the species identifications predicted by the LDA.

#### Evidence of Natural Transformation

In addition to taking measurements of all glochidia found on wild-caught Lake Sturgeon specimens, we also re-examined the hundreds of images taken during this work to identify evidence of larval transformation to the juvenile stage. Here we were looking for the disappearance of the larval adductor muscle, formation of a foot and primordial gill buds, and possibly the appearance of two juvenile adductor muscles.

## Results

### Fieldwork

Voucher specimens and tissue samples are accessioned and catalogued at the Canadian Museum of Nature in the Fish Collection and the National Biodiversity Cryobank of Canada (NBCC), respectively (Table S6).

#### Ottawa River sampling

In total, 1,360 fishes from ∼27 species (some identified only to “sp.”) were collected in the Lac Coulonge region of the Ottawa River in September and November 2021, November 2022, and October-November 2023, including 65 Lake Sturgeon (Table 1). Of these specimens, all 65 Lake Sturgeon and 316 individuals of other species (381 total) were examined for glochidia in the field and/or lab. Fin clips were taken from most specimens collected (Table S6). Of these specimens, we confirmed glochidia on 19 specimens from five species: Lake Sturgeon (n=3), Burbot (*Lota lota*; n=1), Yellow Perch (*Perca ffavescens*; n=3), Sauger (*Sander canadensis*; n=5), Walleye (*Sander vitreus*; n=5).

**Table 1.** Fishes collected in the Lac Coulonge region of the Ottawa River (2021-2023).

| Species | Number collected | Number examined for glochidia | Number with lab-validated glochidia |
|---|---|---|---|
| *Acipenser fulvescens* | 65 | 65 | 3 |
| *Ambloplites rupestris* | 22 | 19 | 0 |
| *Ameiurus nebulosus* | 1 | 1 | 0 |
| *Ameiurus* sp. | 1 | 0 | N/A |
| *Catostomus commersonii* | 40 | 9 | 0 |
| *Corgeonus artedii* | 3 | 3 | 0 |
| *Esox americanus* | 1 | 0 | N/A |
| *Esox lucius* | 11 | 3 | 0 |
| *Etheostoma cf olmstedi* | 108 | 17 | 0 |
| *Hiodon tergisus* | 10 | 4 | 0 |
| *Ictalurus punctatus* | 102 | 55 | 0 |
| *Lepomis gibbosus* | 11 | 5 | 0 |
| *Lepomis macrochirus* | 5 | 0 | N/A |
| *Lepomis* sp. | 1 | 1 | 0 |
| Leuciscidae sp. | 111 | 5 | 0 |
| *Lota lota* | 8 | 8 | 1 |
| *Micropterus dolomieu* | 15 | 13 | 0 |
| *Moxostoma anisurum* | 14 | 13 | 0 |
| *Moxostoma macrolepidotum* | 8 | 6 | 0 |
| *Moxostoma* sp. | 1 | 0 | 0 |
| *Notropis atherinoides* | 187 | 7 | 0 |
| *Notropis heterolepis* | 5 | 5 | 0 |
| *Notropis rubellus* | 375 | 3 | 0 |
| *Perca flavescens* | 121 | 32 | 3 |
| *Percina caprodes* | 13 | 8 | 0 |
| *Pimephales notatus* | 1 | 1 | 0 |
| *Pomoxis nigromaculatus* | 5 | 4 | 0 |
| *Sander canadensis* | 40 | 34 | 5 |
| *Sander vitreus* | 34 | 34 | 5 |
| *Sander* sp. | 2 | 0 | N/A |
| *Semotilus corporalis* | 47 | 8 | 0 |
| **TOTAL** | **1368** | **363** | **17** |

#### St. Lawrence River sampling

We examined 726 Lake Sturgeon in June, July, and August 2021 and none showed any evidence of glochidia on the gills (Table 2). We also retained several dead-in-net bycatch specimens of other species and examined them in the lab but found no evidence of glochidia on these fishes (Table 2). Fin clips were taken from 100 Lake Sturgeon in each of June, July, and August 2021 (300 total) and preserved in 95% EtOH in the field and 42 fin clips or muscle samples were taken from the bycatch specimens in the lab (Table S6).

**Table 2.**
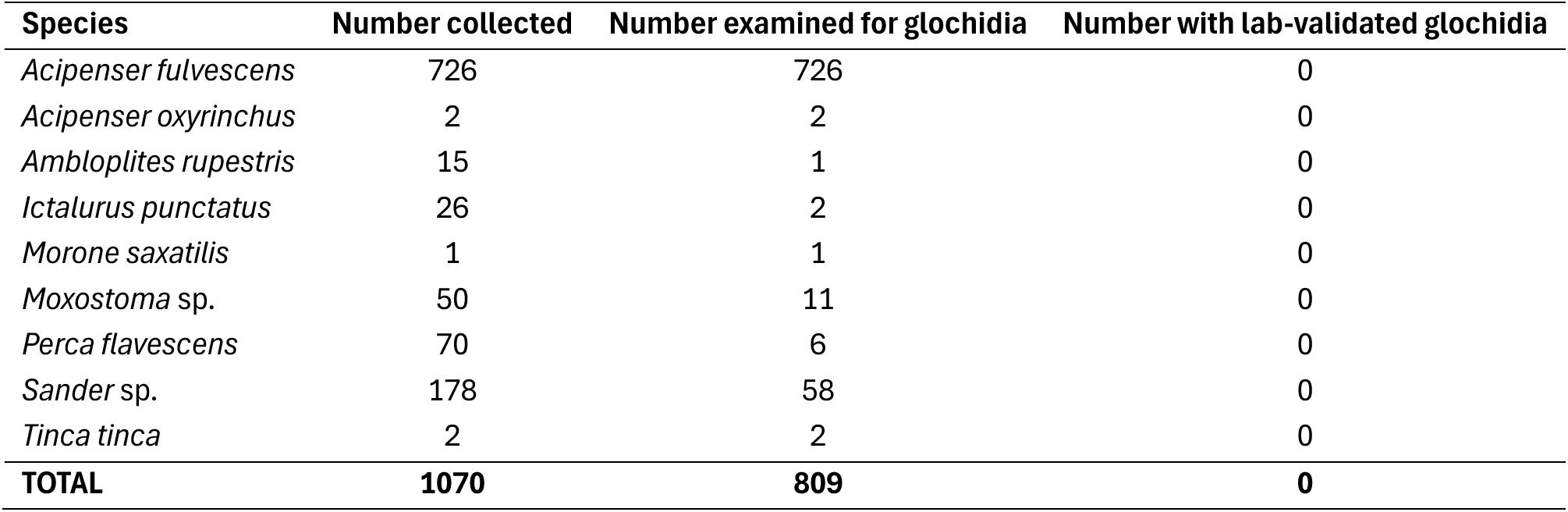
Fishes collected in the Lac Saint-Pierre archipelago region of the St. Lawrence River (2021).

#### Molecular and morphological identification of glochidia

##### Control samples

All COI sequences from the control samples were reciprocally monophyletic by species and clearly distinguishable by COI barcode (Fig. 3, Supp. Mat.). Most species were distinguishable by morphology, particularly if the “whisker” character was available (Fig. 4, Table S7). PC1 explained 91.0% of the variation in the data, with positive PC1 scores associated with decreased shell and hinge length. PC2 explained 7.0% of the variation in the data, with positive PC2 scores associated with increased shell height. PC3 explained very little variation in the data (1.7%), with positive scores being associated with decreased shell depth, but was important for distinguishing Hickorynut morphology from all other species (Fig. S3). Eastern Elliptio, Giant Floater, and Triangle Floater most clearly occupied their own morphospace, while there was considerable overlap across Plain Pocketbook, Eastern Lampmussel, and Black Sandshell. A small number of Hickorynut glochidia showed overlap with some of the Plain Pocketbook glochidia, but Hickorynut glochidia generally had a deeper (wider) shell and, most importantly, displayed a very different hinge whisker structure (Fig. 2).

**Figure 3.**
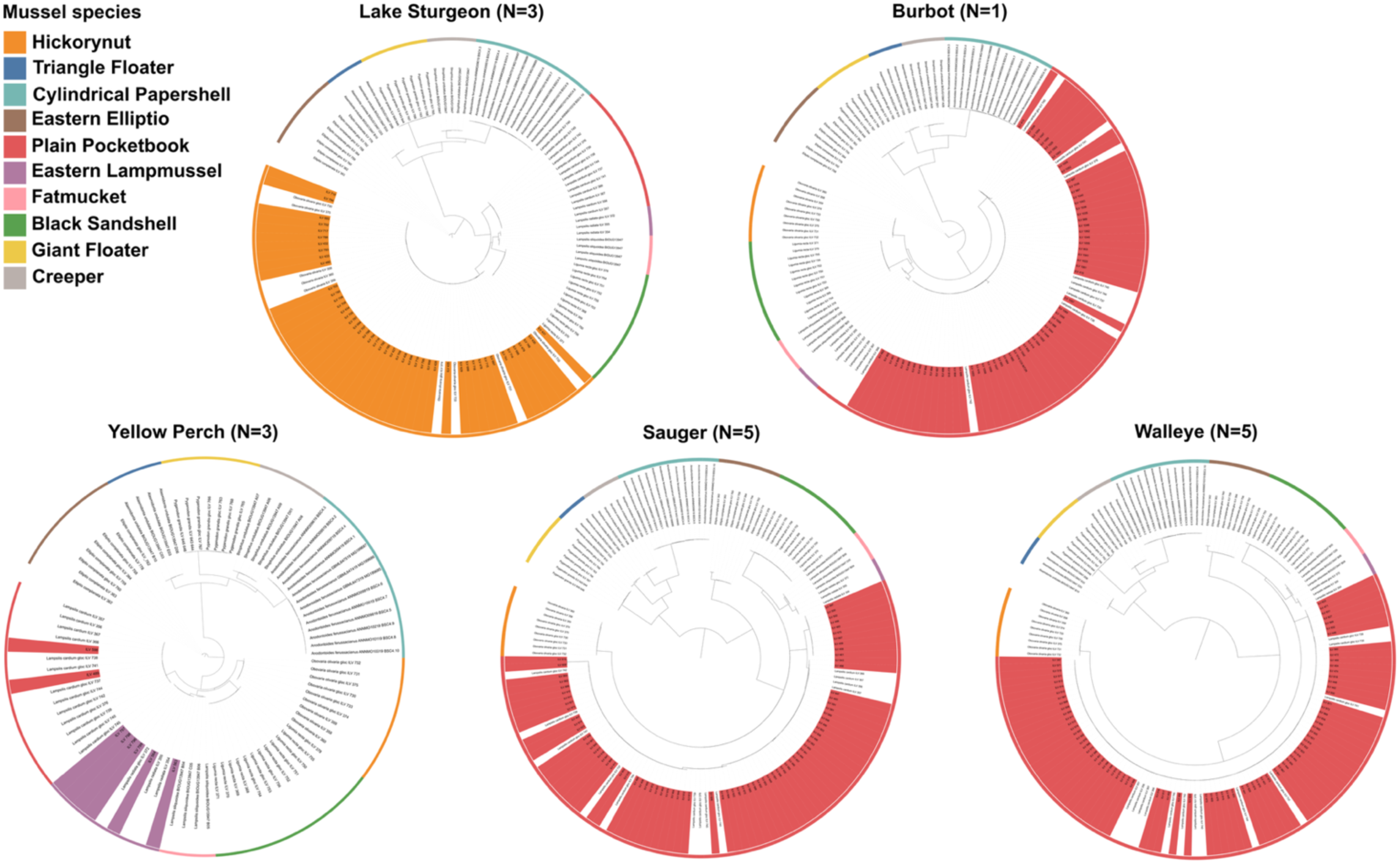
Unrooted bootstrap maximum likelihood (ML) phylogenies of glochidia found on each of the five fish species in the Lac Coulonge region of the Ottawa River with a control dataset of known-identity glochidia and adult mussels. Mussel species are all reciprocally monophyletic with bootstrap support between 97-100%. Tree files are available on Dryad. Figure edited with Inkscape (Inkscape Project (2020) Inkscape (Version 0.92.5). Retrieved from https://inkscape.org)

**Figure 4:**
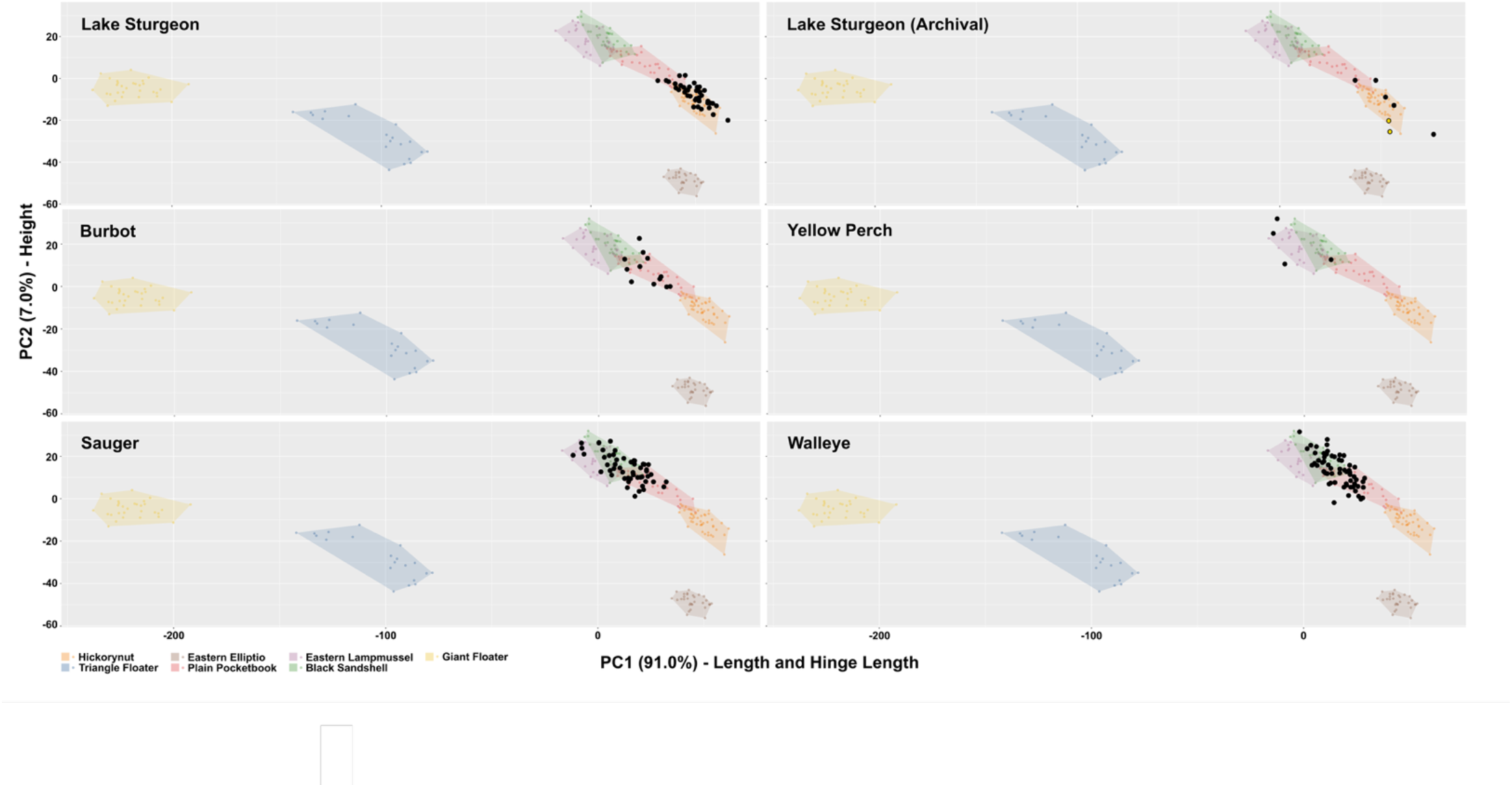
Principal components analysis (PCA) of external glochidia morphology, captured using linear morphometrics. PC1 explains 91.0% of the variation in the data and positive scores are associated with decreases in shell length and hinge length. PC2 explains 7.0% of the variation in the data and positive scores are associated with increase in shell height. Polygons represent the total morphospace occupied by each mussel species, calculated from the stock (control) dataset. Black dots represent the position in morphospace of the glochidia extracted from the gills of fishes in this study. For the Archival Lake Sturgeon panel, black dots represent glochidia extracted from the MELCCPBFP Longueuil specimens, and the yellow dots represent glochidia extracted from the Canadian Museum of Nature specimens. Figure edited with Inkscape (Inkscape Project (2020) Inkscape (Version 0.92.5). Retrieved from https://inkscape.org)

##### Glochidia from wild caught fishes

In total, we obtained COI sequences for 272 glochidia and morphological data for 208 glochidia from the 19 wild-caught fishes from five species. For 146 of these glochidia we had genetic and sufficient morphology data from the same glochidium to assign a species ID using both approaches. Since the statistical analyses required four linear measurements, 35 glochidia with only partial morphological data were excluded, leaving 168 for the statistical and visualization analyses.

The genetic results always produced clear grouping within a species with high bootstrap support (97-100%; Fig. 3, Supp. Data). Given the overlap among several species (Plain Pocketbook, Eastern Lampmussel, Black Sandshell) for glochidial morphology, the morphological identification was sometimes varied and poorly supported (Fig. 4., Table S7). No glochidia extracted from the gills exhibited the presence of hooks, ruling out Triangle Floater and Giant Floater. The presence of whisker type 2 (weak structure) was diagnostic for Hickorynut in this dataset. No glochidia were found with whisker type 1 (no structure), and glochidia with whisker type 3 were ambiguous. For the manual classification based on linear measures, 35 glochidia were labelled “Ambiguous” and 9 labelled “Unknown”. Of the 124 glochidia assigned manually to a single species, 103 glochidia identifications (83.1%) were supported by the LDA predicted identifications. All glochidia classified manually as Eastern Lampmussel (n=4, mean support = 77.4%) and Hickorynut (n=37, mean support = 99.9%) were supported by the LDA. However, for “Ambiguous”, “Unknown”, Plain Pocketbook and Black Sandshell classifications, the LDA predicted identifications were highly variable (Table S7). For this reason, the glochidia found on Burbot, Yellow Perch, Sauger, and Walleye were often manually assigned as “Ambiguous” based on morphology, had incongruence between manual and LDA identifications, and were assigned to a single mussel species with less confidence in the LDA than glochidia found on Lake Sturgeon (Table S7). From COI sequences all glochidia were assigned to species with high support (Fig. 3).

#### Glochidia identity on Lake Sturgeon

All glochidia found on Lake Sturgeon (n=3) were identified as Hickorynut by both genetic (n=46) and morphological (n=48) analyses with strong support (Figs. 3, 4; Tables 3, S7). Although we had morphological measurements for 50 glochidia, two were missing data that precluded species assignment (Table S7). For 43 of the glochidia we had both genetic and morphology data for the same individual glochidium (Table 3). Of the 48 glochidia identified using morphology, 39 were classified using both manual and LDA predictions, while 9 were missing the depth/width measurement and therefore could only be manually categorized, all as Hickorynut (Table S7). Of the 39 with both assignment methods, manual classifications of glochidia were Hickorynut (n=38), and Plain Pocketbook (n=1), while the LDA identified all glochidia as Hickorynut (n=39, average support =99.1%; Table S7). The glochidium with the Plain Pocketbook manual classification did not have a visible hinge characteristic (diagnostic of Hickorynut in this dataset). The three Lake Sturgeon with glochidia were all caught during the second week of November (two in 2021 and one in 2022) and were relatively small (270 mm, 431 mm, and 490 mm TL; Table S6; Fig. S5) compared to the average TL of Lake Sturgeon specimens collected (638.7 ± 138.7 mm).

**Table 3.** Number of individual glochidia with morphology data, DNA data, and both morphology and DNA data. Glochidia lacking some linear measurements and a hinge characteristic could not be assigned to species using either the manual or LDA assignment method. Additional details include Canadian Museum of Nature (CMN) Fish Collection Catalog Number, National Biodiversity Cryobank of Canada (NBCC) or other specimen identifier, and collection date. Species identity column based on genetic identification, unless DNA data unavailable (*).

| Species | CMN Fish Collection Catalog Number | NBCC or other fish specimen identifier | Collection Date | Number of glochidia with some morphology data (number with species assignment) | Number of glochidia with DNA data | Number of glochidia with both morphology & DNA data, with species assignment | Species identity of glochidia |
|---|---|---|---|---|---|---|---|
| *Acipenser fulvescens* | CMNFI 2026-00010.1 | NBCC1002980 | 2021-11-09 | 14 (14) | 11 | 11 | *Obovaria olivaria* |
| *Acipenser fulvescens* | CMNFI 2026-0008.1 | NBCC1002988 | 2021-11-09 | 19 (18) | 18 | 16 | *Obovaria olivaria* |
| *Acipenser fulvescens* | CMNFI 2026-0019.1 | NBCC1003002 | 2022-11-10 | 17 (16) | 17 | 16 | *Obovaria olivaria* |
| *Lota lota* | CMNFI 2026-0018.1 | NBCC1009296 | 2022-11-10 | 17 (17) | 63 | 15 | *Lampsilis cardium* |
| *Perca flavescens* | CMNFI 2026-0007.5 | LC12-1 | 2021-11-08 | 2 (2) | 3 | 2 | *Lampsilis cardium* (1), *Lampsilis radiata* (2) |
| *Perca flavescens* | CMNFI 2026-0007.5 | LC12-3 | 2021-11-08 | 4(4) | 4 | 4 | *Lampsilis radiata* |
| *Perca flavescens* | CMNFI 2026-0012.1 | NBCC1022218 | 2022-11-08 | 0 | 1 | 0 | *Lampsilis cardium* |
| *Sander canadensis* | CMNFI 2026-0009.1 | NBCC1003120 | 2021-11-09 | 12 (12) | 3 | 3 | *Lampsilis cardium* |
| *Sander canadensis* | CMNFI 2026-0009.1 | NBCC1003121 | 2021-11-09 | 12 (12) | 0 | 0 | Ambiguous; highest support for *Lampsilis cardium*\* |
| *Sander canadensis* | CMNFI 2026-0014.1 | NBCC1003126 | 2022-11-08 | 11 (11) | 33 | 11 | *Lampsilis cardium* |
| *Sander canadensis* | CMNFI 2026-0017.2 | NBCC1003140 | 2022-11-10 | 12 (11) | 26 | 8 | *Lampsilis cardium* |
| *Sander canadensis* | CMNFI 2026-0027.2 | NBCC1009093 | 2023-11-02 | 12 (11) | 9 | 8 | *Lampsilis cardium* |
| *Sander canadensis* | CMNFI 2026-0019.2 | NBCC1009303 | 2022-11-10 | 2 (2) | 5 | 1 | *Lampsilis cardium* |
| Sander canadensis | CMNFI 2026-0027.2 | NBCC1030200 | 2023-11-02 | 10 (10) | 0 | 0 | Ambiguous; highest support for *Lampsilis cardium*\* |
| Sander vitreus | CMNFI 2026-0002.1 | NBCC1003056 | 2021-09-21 | 12 (12) | 32 | 12 | *Lampsilis cardium* |
| Sander vitreus | CMNFI 2026-0011.2 | NBCC1003098 | 2021-11-09 | 12 (12) | 2 | 2 | *Lampsilis cardium* |
| Sander vitreus | CMNFI 2026-0016.1 | NBCC1003133 | 2022-11-09 | 18 (18) | 18 | 18 | *Lampsilis cardium* |
| Sander vitreus | CMNFI 2026-0024.2 | NBCC1009087 | 2023-11-01 | 10 (9) | 16 | 9 | *Lampsilis cardium* |
| Sander vitreus | CMNFI 2026-0001.6 | SV1 | 2021-09-20 | 12 (12) | 11 | 10 | *Lampsilis cardium* |

#### Glochidia identity on Burbot

All 63 glochidia that were sequenced for COI from the single Burbot found with glochidia were identified as Plain Pocketbook (Fig. 3) with strong (100%) bootstrap support. Of the 17 glochidia analyzed for morphology, 12 had all four linear measurements and were categorized by both manual and LDA analyses. For the five with manual-only assignment, the identifications were “Ambiguous” (n=4) and Plain Pocketbook (n=1). When both manual and LDA assignment was available, manual assignments resulted in “Ambiguous” (n=1), “Unknown” (n=1), and Plain Pocketbook (n=10), while the LDA identified glochidia as Hickorynut (n=1), Black Sandshell (n=2), and Plain Pocketbook (n=9), with an average support of 91.0% across all identifications (Table S7). The glochidium identified as Hickorynut by the LDA had a whisker type 3, and therefore we are confident it was not a Hickorynut based on morphology. We had genetic data for 15 of these 17 glochidia examined for morphology and all clustered with Plain Pocketbook, including the one identified by LDA as Hickorynut.

#### Glochidia identity on Yellow Perch

Of all the species on which we found glochidia, the Yellow Perch (n=3) had the fewest glochidia (n=8), but was the only fish species where, based on COI barcodes, we identified glochidia from more than one species of mussel: Plain Pocketbook and Eastern Lampmussel. One specimen (LC12-1) with three glochidia had both species present — two Plain Pocketbook and one Eastern Lampmussel; another specimen (LC12-3) collected at the same time from the same site had four glochidia, all of which were Eastern Lampmussel; and the third specimen (NBCC1022218) from a different but nearby site had a single Plain Pocketbook glochidium (Fig. 3, Table 3, Table S7). All these specimens were collected by seining in shallow (<2 m) water. Six of the glochidia were available for morphological analysis, with four being classified using both manual and LDA predictions, while two were missing the depth/width measurement and therefore could only be manually categorized. Unfortunately, the two glochidia identified as Plain Pocketbook by the genetic analysis were unavailable for morphological examination. The two glochidia that had only manual assignments were ambiguous among Eastern Lampmussel, Plain Pocketbook, and Black Sandshell (Table S7). There were conflicting identifications between the manual and LDA identifications for the remaining four glochidia: the LDA identified glochidia as either Eastern Lampmussel (n=2), and Black Sandshell (n=2) with an average support of 62.3% across all identifications, while manual classifications were for Eastern Lampmussel (n=2) or “Ambiguous” (n=2).

#### Glochidia identity on Sauger & Walleye

All 76 glochidia from Sauger (n=5) and 79 glochidia from Walleye (n=5) that amplified and sequenced for COI were identified genetically as Plain Pocketbook (Fig. 3, Table 3, Table S7). Over 60 glochidia for each species were also examined for morphology (Table 3), with species assignment typically as Plain Pocketbook, but also with low support values for Eastern Lampmussel and Black Sandshell (Table S7). Several Sauger were frozen prior to excision of gills, and we presume that the freeze-thaw process damaged the DNA of the glochidia as we were unsuccessful in amplifying COI from glochidia from several of these fishes, despite having intact glochidia for morphological analysis (Tables 3, S7). For the 69 glochidia from Sauger with enough morphology data for species assignment, 17 were missing at least one linear measurement and thus could only be categorized manually as “Ambiguous” (n=10), Plain Pocketbook (n=6) and Eastern Lampmussel (n=1). For the remaining 52 glochidia where both manual and LDA categorization were possible, manual categorization resulted in Ambiguous (n=19), Plain Pocketbook (n=26), Eastern Lampmussel (n=2), and Black Sandshell (n=5) and the predicted identifications from the LDA were Plain Pocketbook (n=30), Eastern Lampmussel (n=7), and Black Sandshell (n=15) with an average support of 78.1% across all LDA identifications (Table S7). For Walleye, only three of the 63 glochidia with sufficient data for assignment were missing the depth/width measurement for LDA analysis, and these were manually categorized as “Ambiguous” (n=1), Plain Pocketbook (n=1), and Black Sandshell (n=1). The remaining 60 were manually categorized as “Ambiguous” (n=14), Unknown (n=5), Plain Pocketbook (n=36), and Black Sandshell (n=5), while the predicted identifications from the LDA were Plain Pocketbook (n=35), Eastern Lampmussel (n=4), and Black Sandshell (n=21) with an average support of 79.5% across all identifications (Table S7).

#### Glochidia from fishes in museum and provincial archival collections

We examined 53 specimens from 33 lots of Lake Sturgeon from the CMN’s Fish Collection for evidence of glochidia and confirmed glochidia on one specimen from CMNFI 2009-0021.1 (Table S5). Similarly, the 13 specimens examined from seven lots from the MELCCPBFP’s Longueuil facility yielded glochidia from two specimens (lots 8883, 74977; Table S5). We were successful in obtaining morphological data from eight of these glochidia (CMN and MELCCPBFP fish collections combined), and most of them clearly grouped with Hickorynut (Fig. 4). Of the eight glochidia, only one was missing the depth/width measurement so could not be included in the LDA analysis; however, the three measurements available (height, length, hinge length) for that glochidium fell with the range of the Hickorynut. For the remaining seven glochidia, one was manually categorized as “Unknown” and identified as Hickorynut by the LDA prediction (support: 100%), one glochidium was manually categorized as Hickorynut and predicted by the LDA as Plain Pocketbook (support: 79.6%), and five were manually categorized and predicted by the LDA as Hickorynut (average support: 100%). The individual glochidium that was identified as Plain Pocketbook by the LDA prediction had the Type 2 whisker, therefore, we are confident it was a Hickorynut and not a Plain Pocketbook. Attempted amplification of COI from seven of these glochidia from two fishes (5 from MELCCPBFP #74977 and 2 from MELCCPBFP #8883) that appeared to have only been preserved in EtOH (i.e., with no exposure to formaldehyde) was unsuccessful. As with the wild-caught Lake Sturgeon from the Ottawa River with confirmed glochidia, the archival specimens with glochidia were all small (<400 mm; Table S5).

#### Evidence of natural transformation of Hickorynut glochidia on wild-caught Lake Sturgeon

Examination of images of the 50 glochidia found on three Lake Sturgeon specimens provided evidence that 14 were in the process of transforming from the larval to the juvenile stage. In those individuals there were clear anatomical indications that metamorphic processes had occurred, including gill buds and foot lobe (Fig. 5, Fig. S4).

**Figure 5.**
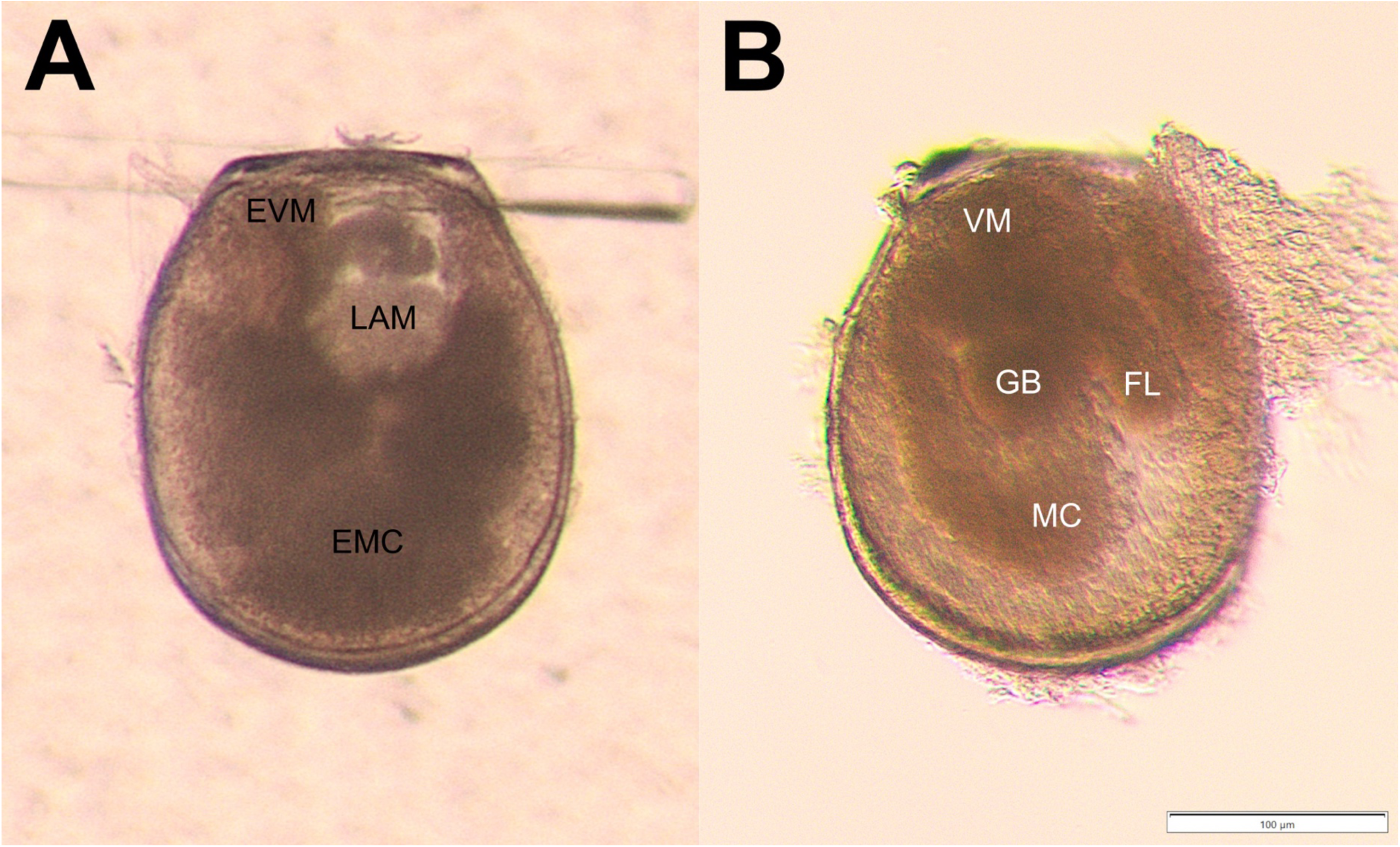
Evidence of Hickorynut (*Obovaria olivaria*) glochidia transformation on wild-caught Lake Sturgeon (*Acipenser fulvescens*). Panel A) Mature glochidium from gravid Hickorynut female. Panel B) Transforming glochidium from Lake Sturgeon (CMNFI 2026-0008.1, NBCC1002988). EVM: embryonic visceral mass, LAM: larval adductor muscle, EMC: embryonic mantle cells, VM: visceral mass, GB: gill bud, FL: foot lobe, MC: mantle cells. Figure edited with Inkscape (Inkscape Project (2020) Inkscape (Version 0.92.5). Retrieved from https://inkscape.org)

## Discussion

### Host fish of Hickorynut in Canada

We confirmed that Lake Sturgeon is a host fish of Hickorynut in Canada with evidence of both natural infestation and natural transformation in fish caught from the Lac Coulonge region of the Ottawa River. Of the ∼30 fish species and 1,172 individuals examined from both the Ottawa and St. Lawrence Rivers, we did not find Hickorynut glochidia on any other species, nor did we find any other species of mussel glochidia on Lake Sturgeon (Figs. 3, 4, 5, S5, Table S7). These results further support speculation that the Hickorynut is a host specialist species and that the Lake Sturgeon is not a host fish for many of the generalist freshwater mussels in Canada.

Of the 27 species of Holarctic sturgeons (Acipenseridae), six species have been identified as host fishes for seven freshwater mussels (Freshwater Mussel Host Database 2017).

According to unpublished data cited by Bouvier et al. (2013), laboratory experiments showed transformation (LT) of numerous Hickorynut glochidia on Shovelnose Sturgeon (*Scaphirhynchus platorynchus*), Pallid Sturgeon (*Scaphirhynchus albus*), and Lake Sturgeon, while other fish species tested (Brook Stickleback (*Culea inconstans*), Central Mudminnow (*Umbra limi*), Mosquitofish (*Gambusia affinis*), Guppy (*Poecilia reticulata*), and Goldeye (*Hiodon alosoides*)) yielded few juveniles. There is also evidence that the invasive Round Goby (*Neogobius melanostomus*) may act as a glochidial sink for Hickorynut and other native mussel species (Tremblay et al. 2016). Atlantic Sturgeon and Freshwater Drum (*Aplodinotus grunniens)* have been suggested as possible Hickorynut hosts (COSEWIC 2011; Clarke 1981, respectively), but there is no evidence from laboratory or field studies for either relationship. Of these species, we only collected two Atlantic Sturgeon and did not find any evidence of glochidia on those specimens (Table 2).

Laboratory experiments with mussels from the southeastern United States have demonstrated that in addition to Hickorynut, Lake Sturgeon are a physiologically suitable host (LT) for Purple Bankclimber (*Elliptoideus sloatianus*; Fritts et al. 2012), but not for five other mussel species (zero transformation [ZT] on: Altamaha Arcmussel (*Alasmidonta arcula*), Altamaha Slabshell (*Elliptio hopetonensis*), Altamaha Lance (*Elliptio shepardiana*), Altamaha Spinymussel (*Elliptio spinosa*), and Altamaha Pocketbook (*Lampsilis dolabraeformis*); Johnson et al. 2012). Note that Lake Sturgeon is not co-distributed with any of these mussel species tested in the laboratory apart from Hickorynut (Fritts et al. 2012; Johnson et al. 2012).

To our knowledge the present study is the first evidence of both natural infestation and natural transformation of Hickorynut glochidia on Lake Sturgeon from any location. This definitive link between two at-risk species can be used to further bolster conservation efforts for each in Canada (e.g., COSEWIC 2011, 2017). Furthermore, through examination of archival fish collections at the Canadian Museum of Nature and a provincial fish collection (MELCCPBFP, Longueuil), we detected measurable and identifiable Hickorynut glochidia on Lake Sturgeon that had been preserved for over 80 years. To our knowledge, this represents the first confirmation of identifiable glochidia from historical collections.

### Utility of DNA barcoding and morphological analyses in glochidia identification

This study employed a joint morphology and genetics approach to the identification of single glochidia to verify the morphology-based methods for distinguishing species and allow for the use of archival fish collections, which are typically unsuitable for genetic analysis, especially for starting material as small as a glochidium. Our glochidia morphology dataset included the three ‘standard’ linear measurements commonly used to compare glochidia species (length, height, hinge length). We have included in the present study a fourth linear dimension (depth/width) as well as a hinge characteristic (‘whisker’ type). Like many others before (e.g., Kennedy and Haag 2005; Tremblay et al. 2015), we found that it was difficult to distinguish some species based on linear measurements (Fig. 4), although the addition of the depth measurement was important for distinguishing Hickorynut glochidia from Plain Pocketbook in particular (Fig. S3) and the hinge characteristic whisker Type 2 was diagnostic of Hickorynut among the species in our dataset (Fig. 2). The inclusion of additional morphological characters (e.g., Klunzinger et al. 2023) will be necessary to distinguish all species from this dataset and the broader freshwater mussel community (ALM et al., in prep).

Studies on in-situ identification of host fishes are becoming more prevalent (e.g., Poelmann et al. 2024; Robinson et al. 2025); however, without bringing live wild-caught fishes into the lab, these studies typically only confirm natural infestation by demonstrating the attachment of mussel glochidia to a host fish, not metamorphosis into a juvenile (natural transformation). Lacking aquaculture facilities to maintain glochidia-infested fishes, we instead used our library of hundreds of images taken for the morphological analyses to look for evidence of transformation of Hickorynut glochidia from Lake Sturgeon (Figs. 5, S4). This approach has been used before as strong evidence for a host relationship between Slippershell Mussel (*Sulcularia viridis*) and Banded Sculpin (*Cottus carolinae*; Zale and Neves 1982).

Our genetic approach for individual glochidia, which used a DNA extraction method developed for forensic science purposes (Rudbeck and Dissing 1998; Malagó et al. 2002) in combination with a two-step (nested) PCR amplification step followed by Sanger sequencing, was highly successful (Fig. 3; Table S6). The segment of COI resulted in reciprocally monophyletic species from our in-house generated control dataset, thus providing unequivocal support for the identification of the “unknown” glochidia species extracted from wild-caught fishes (Fig. 3). This approach is relatively simple and inexpensive and should be widely available and useful to institutions of varying capacities and budgets. A note of caution regarding the preservation of fishes with glochidia for later examination in the laboratory: although we aimed to extract gills with glochidia in the field for immediate preservation in 95% EtOH, timing and conditions (i.e., loss of daylight during November sampling) sometimes required us to freeze fishes that were too large to preserve whole in EtOH. We found that many glochidia from some of these fishes, in particular Sauger (CMNFI 2026-0019.2 [NBCC1009303], CMNFI 2026-0027.2 [NBCC1030200]), did not amplify for COI despite multiple attempts under varying PCR conditions, suggesting that freezing and thawing degraded the DNA (Table 3; Table S7). We recommend immediate preservation of extracted gills in 95% EtOH after dissection in the field, followed soon after by storage at -20 °C or below, whenever possible to avoid potential loss of material for genetic analysis.

### Importance of natural history museum and other archival fish collections in mussel-host fish relationships

The importance of natural history museums for documenting and understanding the origins (evolution), relationships (systematics, ecology), and conservation of biodiversity is well-reported (e.g., Bakker et al. 2020; Hilton et al. 2021; Fedarick et al. 2025; Huang et al. 2025), if not yet fully and widely appreciated. These “libraries of biodiversity” hold answers to many of the most pressing questions in species’ evolution, ecology, and conservation, yet to our knowledge they have not previously been used (or at least published on) for examining fish-mussel relationships, and in particular for identifying host fishes for freshwater mussel glochidia. We demonstrate here the utility of these collections for this purpose by confirming the presence of identifiable Hickorynut glochidia on Lake Sturgeon from the St. Lawrence River that were collected and preserved decades ago (up to 80 years; Fig. 4, Table S5). Glochidia from such historical collections can likely only be analyzed using morphology due to the action of preservatives (often formaldehyde) on DNA degradation; our attempts to amplify COI from glochidia from Lake Sturgeon that appeared to have only been preserved in alcohol (fish with white pupils) was unsuccessful. This emphasizes the need to further refine morphological keys and/or analyses when genetic methods are unavailable.

## Future directions

Our confirmation of Lake Sturgeon as a host for Hickorynut includes fishes from the Ottawa (wild-caught specimens) and St. Lawrence (archival CMN and MELCCPBFP specimens) Rivers. Despite the examination of over 700 Lake Sturgeon from the St. Lawrence River collected in this study during June-August 2021 (Table 2), we did not detect any glochidia on any of these fish. The three Lake Sturgeon from the Ottawa River on which we found glochidia (out of 65 examined) were all collected in November (2021, 2022; Table S6) and none of the Lake Sturgeon collected in this region in September 2021 had evidence of glochidia. This also points to the challenges of examining fish gills from live wild-caught fishes while in the field in fish-mussel connectivity studies. This paucity of Hickorynut glochidia on Lake Sturgeon may be due to a mismatch in the timing of glochidial release by female mussels, which may occur more frequently in winter or early spring. Little is known about the reproductive phenology of Hickorynut anywhere in its North American range; Howard and Anson (1922) report fall release and overwintering of Hickorynut glochidia and Haag (2012) cites a pers. comm about spring release. Brooding females with marsupia filled with fully developed glochidia have commonly been observed in September and October in the Ottawa River (ALM, pers. obs.) and gravid females from well-developed to late stage have been observed August through late November in the St. Lawrence River (AP, pers. obs). Future research involving collection of fishes and Hickorynut during winter months in the Ottawa and St. Lawrence Rivers would likely generate additional valuable knowledge about Hickorynut reproductive phenology and host use in these regions. In addition, all Lake Sturgeon on which we confirmed Hickorynut glochidia, both live-caught and archival, were relatively small (< 500 mm), and thus young (<5 years old). This suggests they likely had naïve immune systems with limited previous exposure to glochidia, as is commonly observed in such host fish relationships (Reuling 1919; Dodd et al. 2006; Ieshko et al. 2016). Further work should target young age classes of potential host fishes. Identification of all potential host fishes will require not only additional fish sampling across the range of Hickorynut across seasons and repeated over years, but also detailed phenological study of Hickorynut in Canada.

Fish collections, both natural history museum and government agency, are likely untapped resources for host fish studies. Work remains to be done to determine how the typical processes of preservation affect glochidia preservation, but we confirmed glochidia on specimens that had been preserved for over 80 years, suggesting these collections are vital resources in understanding fish-mussel connections in nature.

## Supporting information

Supplementary Figures S1-S5 & Buffer Recipes

## Acknowledgements

We thank L. Spotswood and his staff at Spotswood Landing for rearranging docks and lending us his garage so we could set up fish gill examination stations and be sheltered during rainstorms. We thank local resident V. Gervais for his generous contributions in the organization of field trips to Lac Coulonge, providing boat transportation and propane heaters for our cold hands during fish gill examinations on snowy days. We are indebted to D. Lapointe, R. Graham, J. Roelans, and C. Spotswood for providing boat transportation and diving platforms. D. Benoit-Zohr, P.-L. Bastien, L. Lavalley, R. Van Schie, and other members and staff of the Algonquins of Pikwàkanagàn, Kebaowek, and Kitigan Zibi Anishinabeg First Nations for assisting with our work at Spotswood Landing. A. Boudreault from DFO assisted in November 2021. SCUBA divers J. Heinerth and M. D’Aguiar were part of our diving team for the collection of freshwater mussels for the glochidia identification key. Fish sampling in the Lac Saint-Pierre archipelago of the St. Lawrence River was conducted by MFFP/MELCCPBFP teams including D. Hatin, C. Côté, V. Boivin, D. Renaud, N. Auclair, K. Dubois, H. Mercille, K. Jacobs, and T. Gariépy. We are also indebted to A. Hamilton for the development and fabrication of specialized tools and vessels to manipulate glochidia at CMN, and to M. Bainbridge for the natrolite rock crystals for glochidia examination. Thanks also to C. Wilson (OMNR) for the suggestion of the alkaline DNA extraction procedure. M-H. Hubert and T. Martin (fishes) and J-M. Gagnon and B. Postaire (mussels) assisted in processing and cataloguing specimens at CMN. The authors thank the numerous biologists, technicians, students, and collaborators who, over several decades, contributed to the collection and preservation of the data and specimens used in this study. We acknowledge the importance of the historical biological collections maintained by the CMN and MFFP/MELCCPBFP, which made the retrospective analysis possible. Special thanks are extended to MFFP/MELCCPBFP staff M. Lamoureux and Y. Paradis for their assistance with archival records and the preparation of historical Lake Sturgeon specimens from the MFFP/MELCCPBFP fish collections facility in Longueuil and to S. Bernatchez for assistance in the lab. We also recognize the contributions of the many individuals who supported this work under the supervision of successive project leaders, including P. Dumont, D. Hatin, and C. Côté. Y. Paradis, C. Côté, and S. Bernatchez are thanked for their comments on a draft of the manuscript. We thank the Ottawa Riverkeeper organization for assisting us with reaching out to local Ottawa River volunteers and for their support and public outreach on this project. This paper is dedicated to the memory of Vincent Gervais, a friend and incredible volunteer who loved the Ottawa River.

## Author contributions

Conceptualization: ALM, KLI, H-FE, TH, NRA, AP

Data curation: KLI, ALM, SES

Formal analysis: KLI, SES

Funding acquisition: H-FE, ALM, KLI

Investigation: ALM & KLI, TH, SES, NRA, AL, EM, RDB, SF, BRA, ST, AP, M-HF, H-FE

Methodology: KLI & ALM, SES, RDB, SF, TH

Project Administration: ALM & KLI, TH, H-FE, SES

Resources: KLI, ALM, TH, SES, RDB, ST, AP, M-HF, H-FE

Supervision: KLI & ALM, TH, H-FE

Visualization: SES, KLI, ALM, BRA

Writing – original draft: KLI

Writing – review & editing: KLI, SES, ALM, BRA, TH, NRA, AL, EM, RDB, SF, ST, AP, M-HF, H-FE

## Author information

### Data availability

Data generated or analyzed during this study are provided within the published article and its Supplementary Materials (figures, tables, buffer recipes), Supplementary Data (Dryad - phylogenetic tree files; https://doi.org/10.5061/dryad.hhmgqnkzn), and The Barcode of Life Data System (BOLD) data portal (COI barcode sequences; BOLD Process IDs OOAC001-26 – OOAC324-26).

