## Supplementary Figures S1-S5 & Buffer Recipes for "Validation of the host fish relationship between Hickorynut (*Obovaria olivaria*) and Lake Sturgeon (*Acipenser fulvescens*) from wild-caught and archival museum fishes"

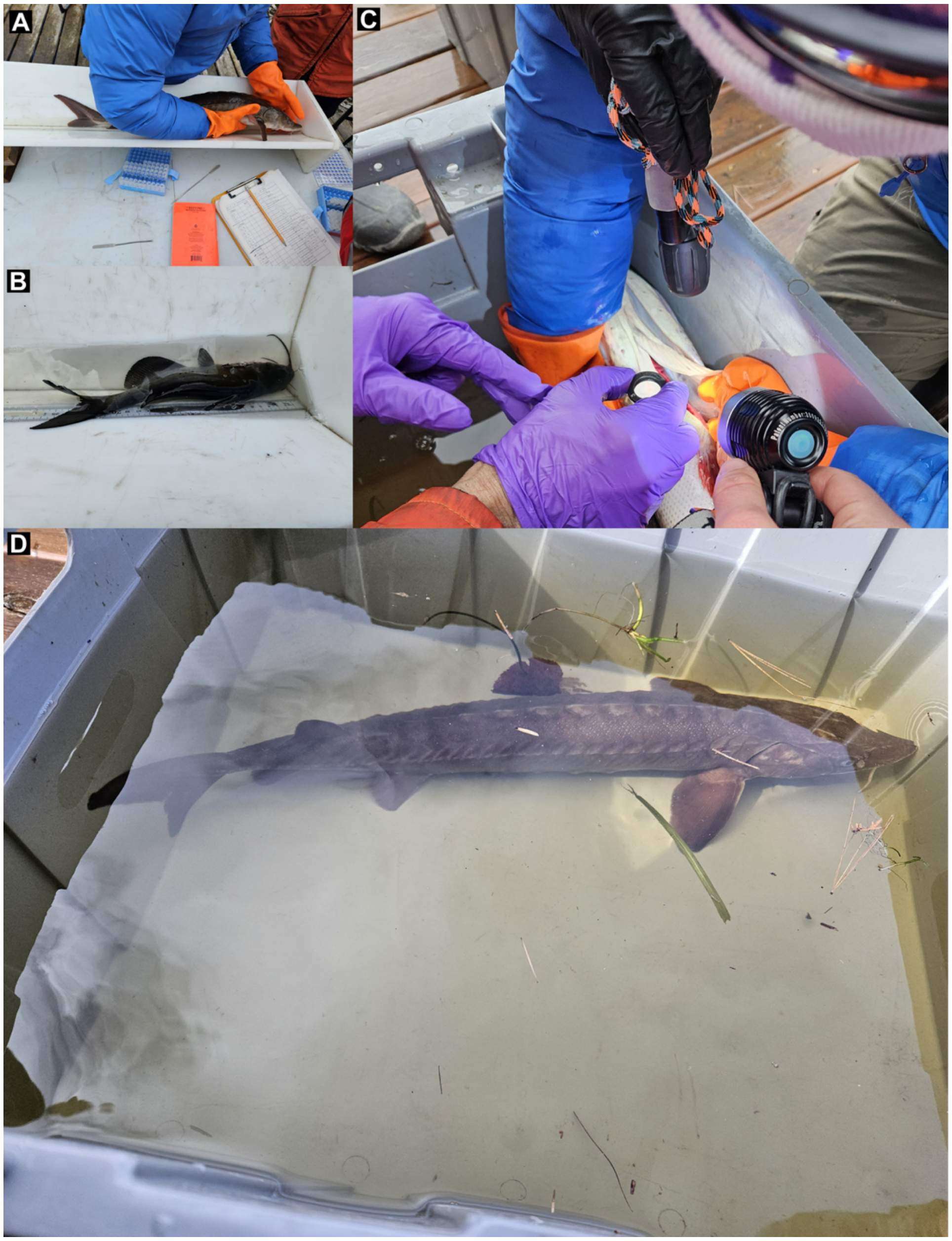


Figure S1. Field set-up for examining live fishes for glochidia. A & B. Careful placement of fish into measuring trough with river water to immerse gills on side not being examined. C. Examination of live fish for evidence of glochidia using hand lens. D. Recovery bin for fish prior to release back into river.

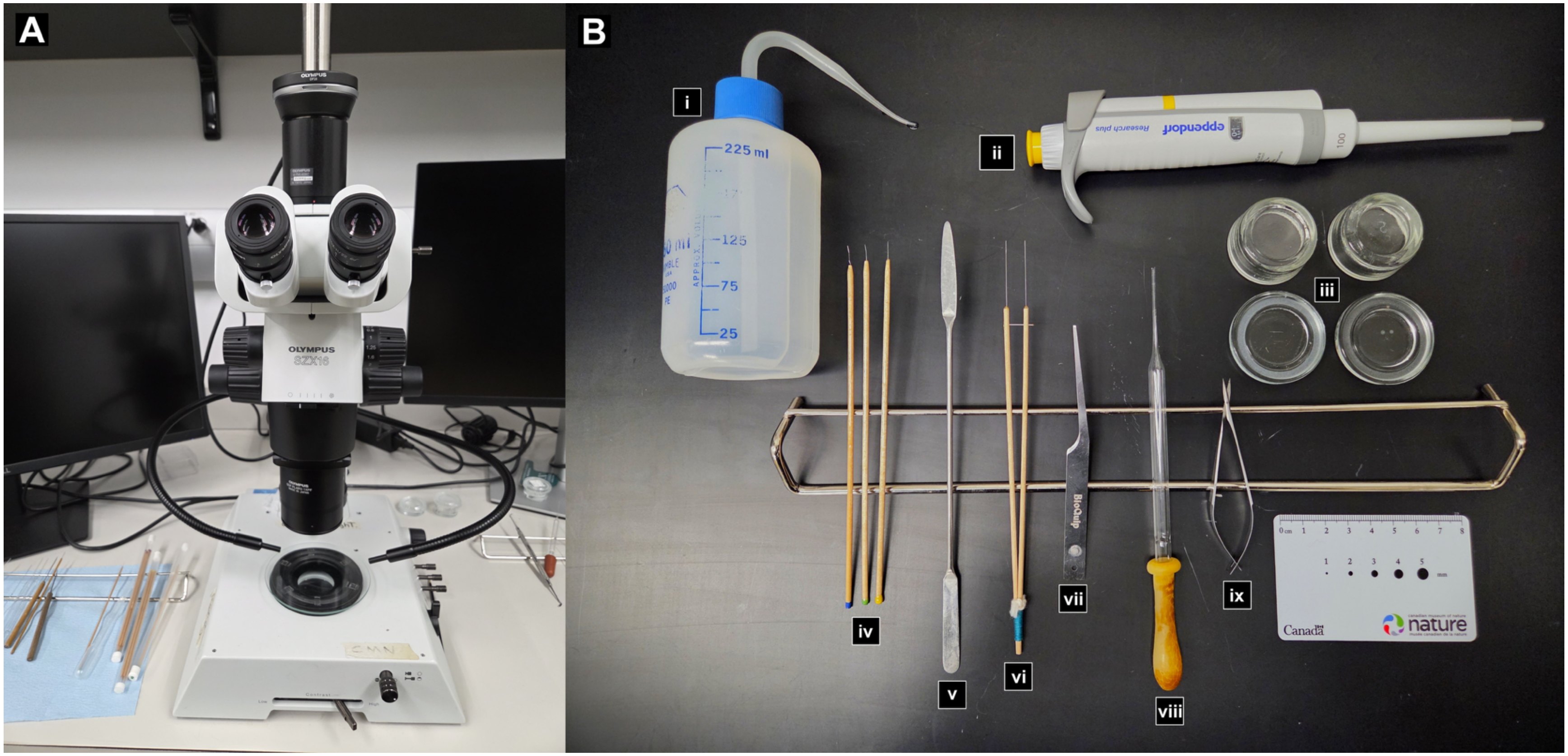


Figure S2. Images of custom laboratory set-up for measuring and imaging individual glochidia. A. Olympus SZX16 stereo microscope. B. Various tools and supplies. i) squirt bottle with 30% ethanol, ii) micropipette, iii) glass vessels with small pits, iv) microneedles, v) spatula, vi) custom forceps, vii) pliable forceps, vii) glass pipette, ix) microscissors. Specialised tools for glochidia manipulation, including glass dishes with micro-wells and troughs, designed by A. Hamilton and A.L. Martel, and built by A. Hamilton.

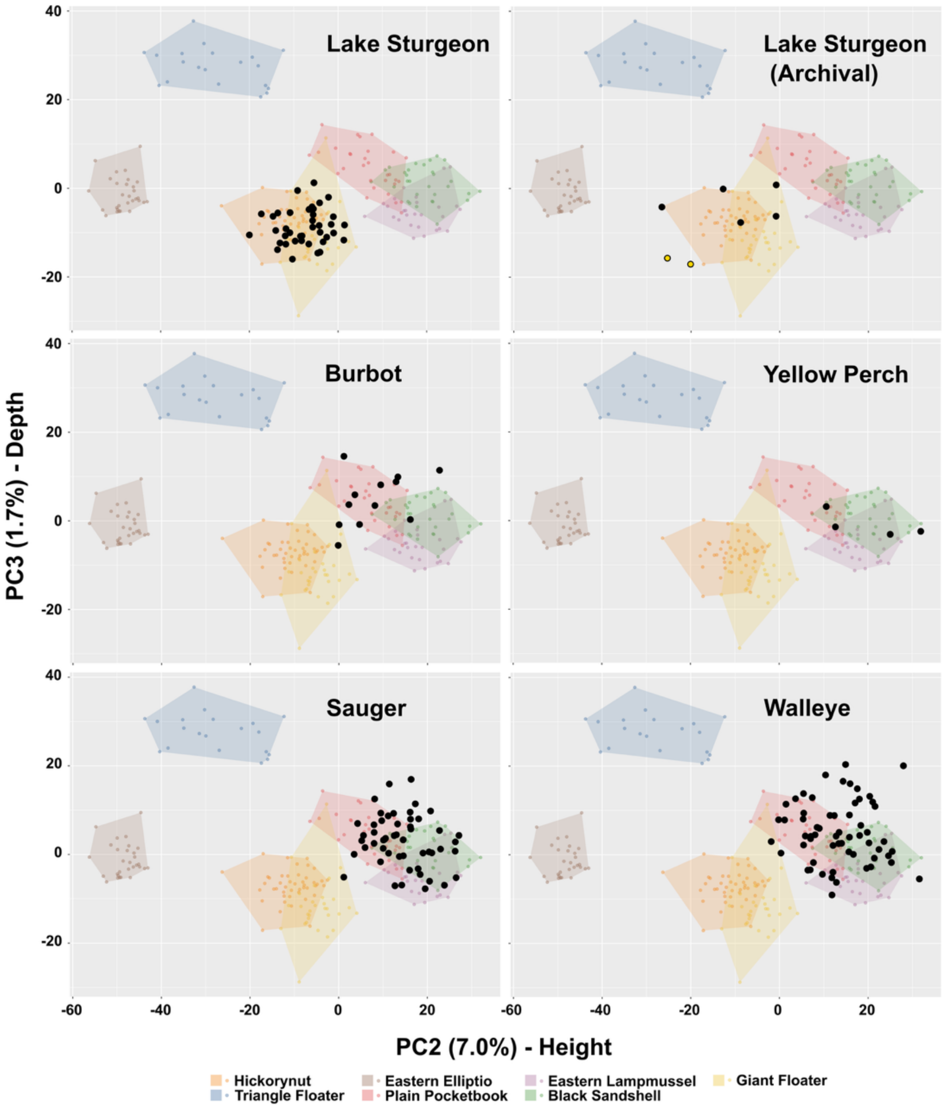


Figure S3. Principal components analysis of external glochidia morphology, captured using linear morphometrics. PC2 explains 7.0% of the variation in the data and positive scores are associated with increase in shell height. PC3 explains 1.7% of variation in the data and positive values are associated with decreased shell depth. However, PC3 was important for distinguishing Hickorynut morphology from all other species. Polygons represent the total morphospace occupied by each mussel species, calculated from the stock (control) dataset. Black dots represent the position in morphospace of the glochidia extracted from the gills of fishes in this study. For the Archival Lake Sturgeon panel, black dots represent glochidia extracted from the Ministère de l'Environnement, de la Lutte contre les changements climatiques, de la Protection de la Biodiversité, de la Faune et des Parcs (MELCCPBFP) Longueuil specimens, and the yellow dots represent glochidia extracted from the Canadian Museum of Nature (CMN) specimens.

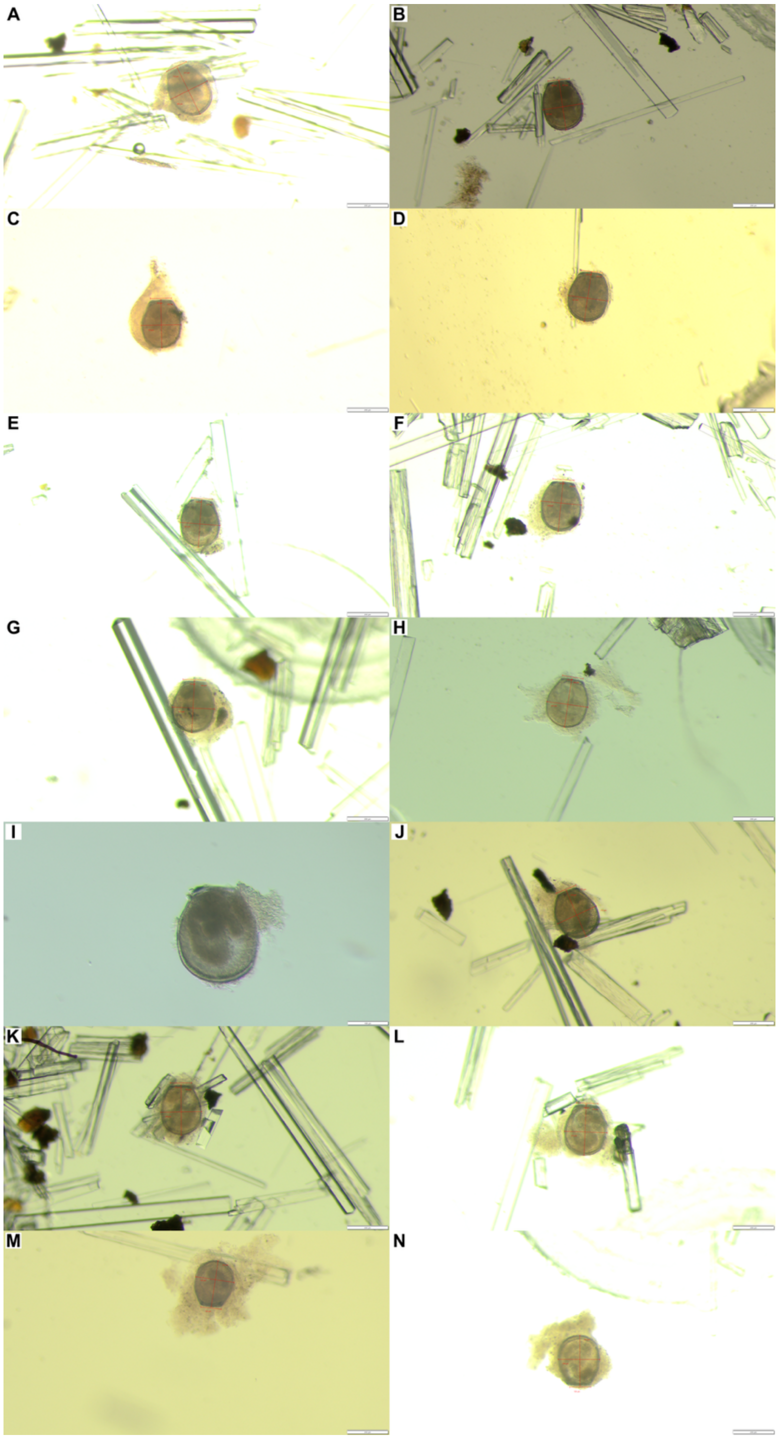


Figure S4. Images of 14 Hickorynut (*Obovaria olivaria*) glochidia from wild-caught Lake Sturgeon (*Acipenser fulvescens*) that show evidence of natural transformation from the larval to the juvenile stage. A-F from CMNFI 2026-00010.1 (NBCC1002980), G-N from CMNFI 2026-0008.1 (NBCC1002988).

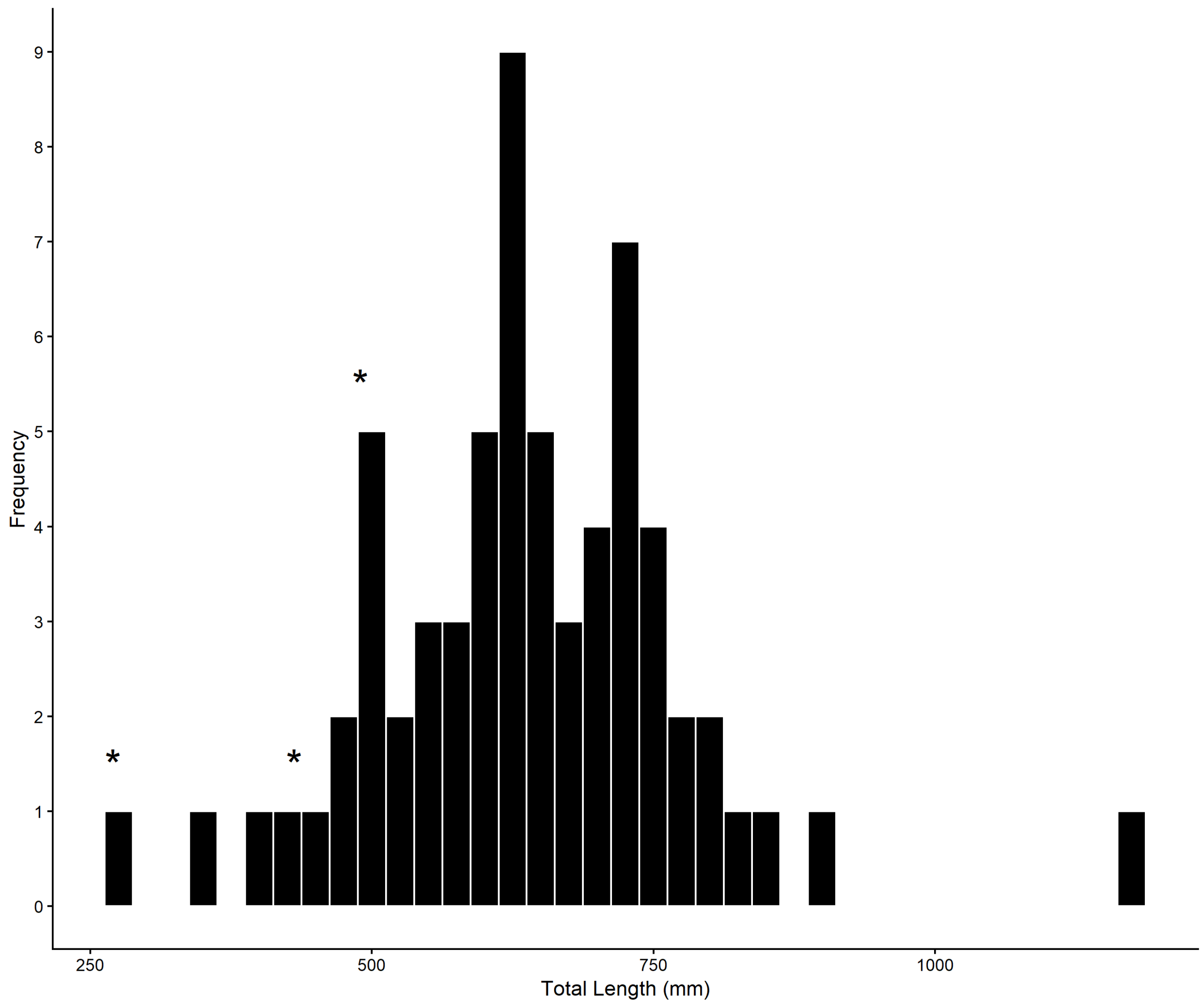


Figure S5. Length frequency histogram of Lake Sturgeon collected in the Lac Coulonge regions of the Ottawa River and examined for glochidia.

### DNA Extraction Buffer Recipes

#### AL Buffer (Animal Lysis Buffer)

| Reagent | Stock conc'n | For 200 ml | Final conc'n |
|---|---|---|---|
| SDS | | 1.0 g | 0.5% |
| NaCl | 5 M | 4 ml | 100 mM |
| Tris-HCl | 2 M, pH 8 | 5 ml | 50 mM |
| EDTA | 0.5 M, pH 8 | 4 ml | 10 mM |
| $ddH_2O$ | | 187 ml | |

Place a 200 ml autoclavable storage bottle onto a balance, tare, and add the SDS.

Remove the bottle from the balance and add NaCl, Tris-HCl, EDTA and $ddH_2O$. Place bottle on a stir plate, add a magnetic stirrer, and stir until SDS is fully dissolved.

Twist lid onto bottle loosely. Autoclave at 121°C for 20 minutes.

Store at room temperature.

---

#### AMB Buffer (Animal & Macroalgae Binding Buffer)

| Reagent | Stock conc'n | For 200 ml | Final conc'n |
|---|---|---|---|
| GuSCN (MW = 118.2 g/mol) | | 141.84 g | 6 M |
| Tris-HCl | 0.1 M, pH 6.4 | 20 ml | 10 mM |
| EDTA | 0.5 M, pH 8 | 8 ml | 20 mM |
| Triton X-100 | | 8 ml | 4% |
| $ddH_2O$ | | to 200 ml | |

Place a beaker onto a balance, tare, and add the GuSCN.

Remove beaker from the balance and add remaining ingredients. Place on a stir plate set to 100°C and leave to stir until the GuSCN is fully dissolved.

Decant the buffer into a 200 ml graduated cylinder and add $ddH_2O$ to a total volume of 200 ml if needed.

Store at room temperature.

If any re-crystallization occurs, pre-warm at 56°C to dissolve before use.

#### Tris-HCl, pH 8, 2 M (tris hydroxymethylaminomethane, $NH_2C(CH_2OH)_3$, and hydrochloric acid)

| Reagent | For 200 ml |
|---|---|
| $ddH_2O$ | approx. 140 ml |
| Tris base (MW = 121.14 g/mol) | 48.46 g |
| HCl | Add with pipette until pH = 8.0 |

Add the $ddH_2O$ and the Tris base to a beaker and mix on a stir plate.

Measure the pH with a pH meter and add HCl to achieve the target pH. Start by adding 15 ml of HCl but expect to add about 20 ml to achieve the target pH.

Once pH = 8.0, bring volume close to 200 ml by adding more $ddH_2O$. Recheck pH before setting volume at 200 ml.

---

#### SW Buffer (Second Wash Buffer)

| Reagent | Stock conc'n | For 475 ml | Final conc'n |
|---|---|---|---|
| Ethanol | 99% | 290 ml | 60% |
| NaCl | 5 M | 4.75 ml | 50 mM |
| Tris-HCl | 1 M, pH 7.4 | 4.75 ml | 10 mM |
| EDTA | 0.5 M, pH 8 | 0.475 ml | 0.5 mM |
| $ddH_2O$ | | to 475 ml | |

Add reagents to a 500 ml graduated cylinder.

Decant into a storage bottle and swirl to mix until homogenous.

Store at -20 °C.

---

#### THE Buffer (Tris Half-EDTA Buffer)

| Reagent | Stock conc'n | For 200 ml | Final conc'n |
|---|---|---|---|
| Tris-HCl, pH 8 | 2 M, pH 8 | 1 ml | 10 mM |
| EDTA | 0.5 M, pH 8 | 0.2 ml | 0.5 mM |
| $ddH_2O$ | | 198.8 ml | |

Add reagents to a 200 ml autoclavable storage bottle. Swirl to mix. Twist lid onto bottle loosely. Autoclave at 121 °C for 20 minutes.

Store at 4 °C.
